# Dissecting errors made in response to externally- and internally-driven visual tasks in the common marmosets and humans

**DOI:** 10.1101/2021.08.29.458139

**Authors:** Wajd Amly, Chih-Yang Chen, Hirotaka Onoe, Tadashi Isa

## Abstract

Various visual paradigms in oculomotor research have been used for studying the neural processes of eye movement, cognitive control, attention and neurological disorders. However, we usually analyse data collected from humans or over-trained non-human primates (NHPs), focusing only on successful trials, whereas error trials are usually excluded. These errors may repetitively show up in diseases, but their interpretation would be difficult due to the absence of records taken from healthy controls. In the current study, we aimed to analyse both correctly and incorrectly performed trials in both marmosets and humans. We trained marmosets to perform the gap saccade task and the oculomotor delayed response task. We also collected data from human subjects who performed identical tasks. We categorised error trials into three different groups, based on the time when an incorrect response occurred. We also interpreted possible causes by analysing saccade reaction time, saccade landing position and task history. Despite the rareness of human error, we found that marmosets and humans showed remarkably similar behaviour in error and success. We also found that successful saccades in the gap saccade task had always the highest peak velocity in both species, reflecting faster sensorimotor processing for correct responses. Our results suggest that marmosets and humans might share similar neural processing for successful and unsuccessful oculomotor behaviour, making them a suitable model for studying human behaviour. More importantly, analysing error trials in sync with successful ones will provide further insights into the cognitive and sensorimotor processes.

**NEW AND NOTEWORTHY:** This is the first detailed report focusing on analysing both error and successful trials in oculomotor tasks. We proposed nomenclatures and a generalizable way of grouping and analysing error trials. Our results also indicate that marmosets can be a promising experimental candidate for oculomotor research because they replicate the saccade properties of error and success seen in humans. This will help set the baseline measurements to study brain disorders using NHP and understand the neural mechanisms from a different perspective.

## INTRODUCTION

Saccades are conjugate rapid eye movements that primates make to bring the target of interest to the centre of the fovea (Yarbus, 1967; Weymouth et al. 1928). The measurable reaction time and the stereotypic kinematics of saccades (Bahill et al., 1975; Lebedev et al., 1996; Robinson 1981) makes them a valuable behaviour model for probing neural mechanisms for both basic and clinical research.

Saccadic eye movements were first characterized in awake macaque monkeys by Fuchs and colleagues (Fuchs 1967; Fuchs and Robinson 1966). Since that time, macaques have become the primary model for studying the cortical and subcortical control of saccades (for reviews, see Borra and Luppino 2019; Coiner et al. 2019; Schiller and Tehovnik 2009; Takahashi and Shinoda 2018). The interdependency between the cortical saccadic system and more cognitive phenomena such as attention, visual working memory, and visual decision-making has also been reported (Boshra and Kastner 2022; Hunt et al. 2019; Jonikaitis and Moore 2019; Spering 2022). Saccades are now not only to study the oculomotor system itself, but their use in research has extended to observing and evaluating cognitive processes (Beesley et al. 2019; Hafed et al. 2015; Liversedge et al. 2011; Wolf and Lappe 2021). Macaques, with their ability to learn and perform cognitive tasks with close-to-perfect performance, have simultaneously become a mainstay in neural research. This has led to a situation where the number of error trials is too small to be analysed, and they are often not included in the reported results.

Over recent years, oculomotor paradigms including saccadic behaviour as a marker have been extended to clinical investigations as an early diagnostic behavioural biomarker. They have been used to evaluate different neurological disorders (Gooding and Basso 2008; Das et al. 2022; Everling and Fischer 1998; Leigh and Kennard 2004; Ramat et al. 2007). Such disorders include movement disorders (Termsarasab et al., 2015; Waldthaler et al., 2021; Fukushima et al., 2017), psychiatric diseases (Bittencourt et al. 2013; Morita et al. 2020), and neurodegenerative diseases (Antoniades and Kennard 2015; Opwonya et al. 2021). In those studies, the patient populations make different kinds of eye movement errors while performing behavioural tasks. While patients often exhibit obvious deficits in behaviour, it is not clear how to connect these deficits to the error typically made even by healthy subjects. This inadequate understanding of how errors occur in healthy controls makes it challenging to further diagnose the affected brain structures. Recently, macaques have been used to study neurological disorders, justified by their similar brain structure and function to humans (Hutchison and Everling 2012). Indeed, several studies have successfully used macaques to reproduce the symptoms of various brain disorders (Qiu and Li 2017; Verdier et al. 2015; Sherman et al. 2021; Souder et al. 2021; Liu et al. 2016). However, the primate models were not tested using many of the oculomotor behaviour paradigms commonly used in human patients, making a direct comparison between patients and animal models difficult.

The common marmoset (*Callithrix jacchus*), a new world monkey, has been proposed as an alternative NHP model which is easier to manipulate transgenically (Izpisua Belmonte et al. 2015; Park et al. 2016; Sasaki et al. 2009), and has a shorter lifespan, enabling long-term behavioural follow-up of the progression of neurological diseases (Tardif et al. 2011). Several neurological disease models have also been successfully generated in marmosets using transgenic or other means (Eslamboli et al., 2007; Tomioka et al., 2017; Geula et al. 2002; Philippens et al. 2016; Ridley et al. 2006 Jenner et al., 1984; Fox et al., 2010; Emborg, 2017; Sawamura et al. 2022). Marmosets can perform a variety of visual and cognitive tasks under head unrestrained (Clarke et al. 2011; Jendritza et al. 2021; Spinelli et al. 2004; Yamazaki et al. 2011; Okano 2021) and head restrained conditions (Mitchell et al. 2014; Johnston et al., 2018; Chen et al., 2021; Sedaghat-Nejad et al. 2019), and they exhibit saccadic behaviour similar to humans under oculomotor paradigms (Chen et al. 2021).

However, reports of the similarity of marmoset and human saccadic behaviour compared only on *successful* trials. Given no reason to doubt that error trials would differ, we hypothesise that marmoset saccade properties in error trials are also similar to those of humans. To investigate this, we trained three marmosets in two different visual paradigms, the externally-driven gap saccade task (gap task) and the internally-driven oculomotor delayed response (ODR) task (Funahashi et al. 1989; Hikosaka and Wurtz 1983) concurrently, mimicking clinical studies in human patients. Three human subjects also performed the same tasks. Here, we present results from both successful and unsuccessful (error) trials, and propose a systematic way to categorize error trials. We identify the different types of errors, analyse their possible causes, and compare them between the two species. The overarching goal of this research is to establish the baseline profile of error saccades, thus paving the way for future studies of human neurological disorders using marmosets.

## METHOD

### Ethics Approval

All marmosets (Callithrix jacchus) used in this study were born in the breeding colony of Kyoto University Animal Research Facility. The experiments followed the guidelines of the Japan Neuroscience Society and the Science Council of Japan. The animal Ethics Committee at Kyoto University, Japan approved all the experimental procedures.

Ethics committees at the Medical Faculty of Kyoto University approved human experiments. Subjects provided informed, written consent to join the experiment.

### Animal Preparation and Surgical Procedure

Three adult marmosets participated in this study; marmoset S: 2.5 years old male, 390 g; marmoset N: 2.5 years old female, 370 g; and marmoset M: 2 years old male, 340 g. All the marmosets started with habituation to the transportation cage This was followed by two weeks of chair training to familiarise the marmosets with the primate chair and the lab environment. Later, marmosets underwent headpost implantation surgery to prepare for head fixation. Anaesthesia was induced using 14 mg/kg ketamine and 0.14 mg/kg medetomidine. In addition, 0.07 mg/kg atropine, 0.5 mg/kg dexamethasone, and 1.5 mg/kg gentamicin were administered to prevent excessive salivation, pain and infection respectively. The eyes were protected with erythromycin-containing eye ointment. Under 1.5 % isoflurane, each marmoset was aseptically mounted with its custom-designed headpost based on the MRI reconstruction of the skull. The headposts were 3D-printed using Dental SG Resin (Form2, Formlabs, U.S.A.), and were attached to marmosets’ skulls using Super-Bound (Sun Medical Co., Ltd., Japan). After the surgery, marmosets were given the antibiotic ampicillin, 30 mg/kg, and the anti-inflammatory and analgesic medication ketoprofen, 0.8 mg/kg for 4 days. After the medical treatment, an additional week was given to allow for a full recovery.

### Human Subjects

Three adult human subjects aged 29–37 participated in this study, two males (human C, H) and one female (human W). Humans’ head position was stabilized using a custom-made chin rest.

### Monitor Setup and Visual Stimulus Software

We used a 1280×1024 pixel resolution LCD monitor (Dell P1917S) with a refresh rate of 60 Hz for marmosets and a 1920×1080 pixel resolution LCD monitor (Dell AW2125HF) cropped to 1280×1024 pixel with the same refresh rate for humans. The visual distance was 41 cm, and the stimuli were presented on a grey background with a luminance of 80.81 cd/m^2^ for marmosets, and 58.47 cd/m^2^ for humans. We used PsychoPy 3.6 (Peirce et al. 2019) to control the stimuli display and the subjects’ eye position.

### Eye Tracking and Calibration

An Eyelink 1000 Plus (SR Research, Canada) tracked marmosets’ eyes binocularly with a sampling frequency of 500 Hz. We used both pupil and corneal reflection information for accurate tracking. We calibrated the eyes’ position with the same method described by (Chen et al. 2021). Briefly, for the online calibration, we used marmosets’ face photos (8 images per trial, sizes ranging from 4 to 20.25 deg^2^; distance from the monitor centre to the image centre ±12 deg horizontal and ±10 deg vertical) presented at random eccentricities on the screen and the marmosets were free to look at the photos for 10 seconds per trial. We then examined the presented photos and the corresponding marmosets’ gazes. We estimated the x and y gain and offset of the eye position for each marmoset individually. This calibration was done once for each marmoset after sitting in front of the eye tracker for the first time. We validated the calibration result every day by taking the average eye position 0 to 50 ms before reward delivery from the blocked gap task (described below) and calculated the offsets and gains using simple linear regression. The offset and gain results were updated to the online task controller; a Psychopy-written script to monitor the eye position. An Eyelink 1000 (SR Research, Canada) was used for human subjects, but all the other eye tracker settings and calibration procedures were similar to those used for marmosets.

### Visual Tasks

#### Blocked Gap Task

In the gap task (**Fig. 1A**), target eccentricity (4, 6 or 8 degrees) and gap period (0-400 ms with 50 ms increment) were predetermined randomly before the session onset and fixed for each session following macaque research (Chen et al. 2016; Li et al. 2021; Parẻ and Munoz 2001) and to facilitate the training. The target location was randomly selected from eight different locations equally spaced by a 45-degree radial visual angle every trial.

**Figure 1.**
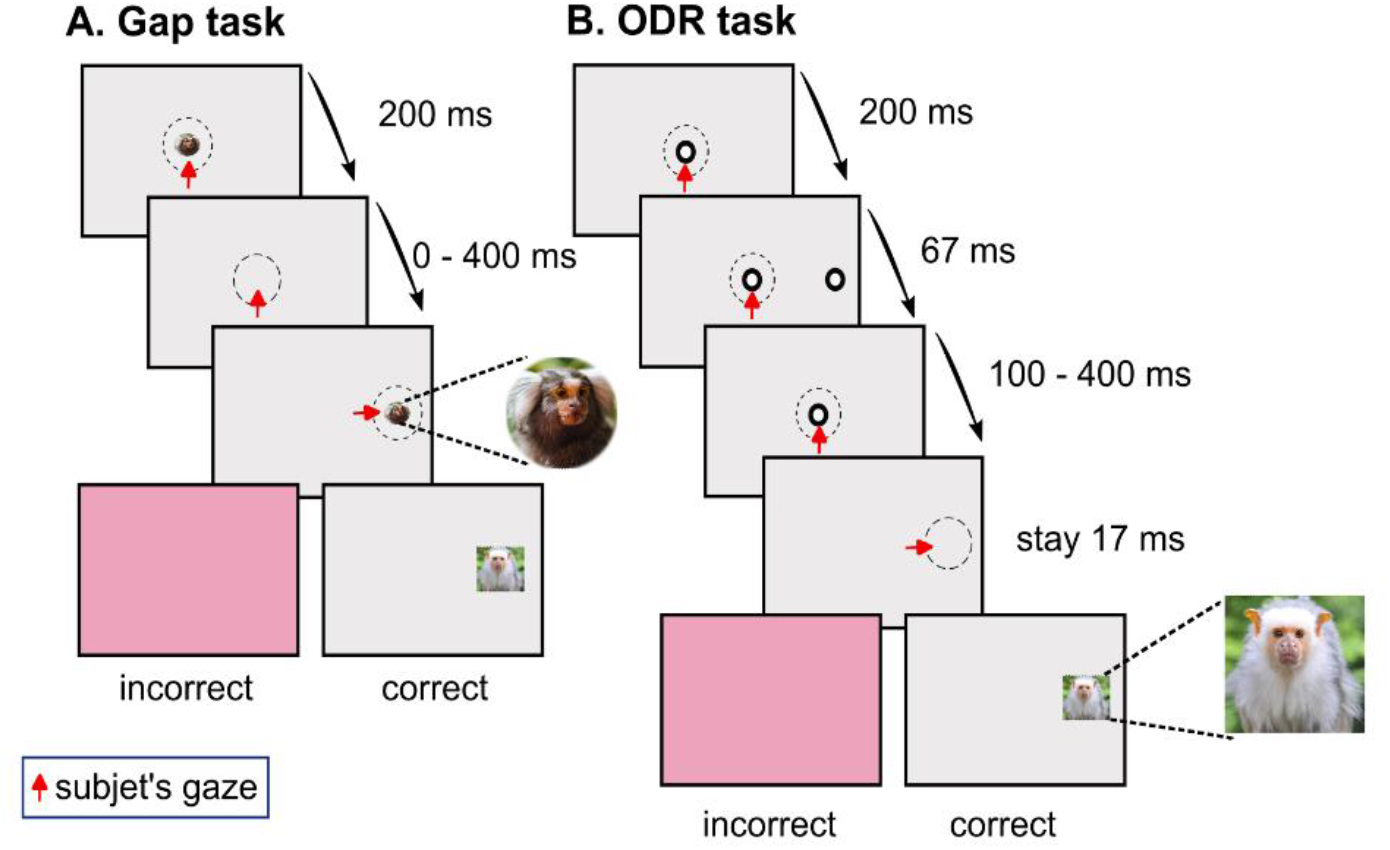
Tasks illustration. **A)** In the gap task, a marmoset face was used as the stimulus. Each trial started with a fixation stimulus that appeared at the screen centre for the subject to fixate for 200 ms. Following the fixation stimulus offset, a gap period (0 - 400 ms with 50 ms increment) followed, and the gaze must be maintained at the centre. At the end of the gap period, a visual target appeared, and the subject must make a saccade towards the target and maintain the gaze for 200 ms. If the answer was correct, a rewarding photo was displayed but if the answer was incorrect, a one-frame red screen was flashed. **B)** In the ODR task, a black circle with a white centre was used as the stimulus. The fixation stimulus appeared at the screen centre for the subject to fixate for 200 ms. A peripheral target flashed for 67 ms after the initial fixation, and the gaze must be maintained at the centre. Then, a delay period (100 - 400 ms with 50 ms increment) followed and the gaze must be still maintained at the centre. At the end of the delay period, the fixation stimulus was turned off and the subject must make a saccade towards the remembered target location and maintain the gaze for around 17 ms. If the answer was correct, a rewarding photo was displayed but if the answer was incorrect, a one-frame red screen was flashed. Dashed circles indicate the eye window which was invisible to the subjects. Refer to **METHOD** for a detailed description.

A trial started by presenting a 1-degree-radius marmoset face as the fixation stimulus at the centre of the monitor over a grey background for up to 1000 ms. The subject had to look and hold the gaze within a 2-degree-radius invisible eye window centred at the fixation stimulus (**Fig. 1A**, dashed circle in the top graph). If the gaze was successfully maintained for 200 ms, the fixation stimulus was removed, and the subject had to maintain a steady fixation during the gap period (**Fig. 1A**, middle graph) until the visual target (marmoset face, go cue) appeared. The subject was then required to make a saccade to the target within 1000 ms and maintain fixation for 200 ms to attain a drop (0.05 ml) of the liquid reward along with a rewarding photo that was displayed at the target location (**Fig. 1A**, bottom graph). The trial was aborted if the subject failed to initiate fixation within the defined time after trial initiation, if the gaze escaped the fixation window before the target onset during fixation or gap periods, if the gaze escaped the target eye window before the time allowed so, or if the subject never looked to the target within the 1000 ms response period. A one-frame red screen flash was presented as an indication of the trial’s failure. One session was terminated if either 24 successful trials (see Result section) were collected or if a maximum of 48 trials (including error) were collected. The marmoset face used for both the fixation and the target varied in each session. In this task, faces were used as stimuli to aid the marmosets in differentiating this task from the oculomotor delayed response task below.

### Blocked Oculomotor Delayed Response Task (ODR)

In the oculomotor delayed response (ODR) task (**Fig. 1B**), target eccentricity (4 or 6 degrees) and delay period (100-400 ms with 50 ms increment, (similar to Andersen et al. 1990; Massendari et al. 2018; Willeke et al. 2019; Angelo V. Colmenero 2021) were predetermined before the session onset. Similar to the gap task, these parameters were fixed for each session following macaque research (Chen et al. 2016; Kagan et al. 2010; Lowe et al. 2022; Tsujimoto and Sawaguchi 2004) and to facilitate the training. We used a different fixation and target stimuli (a black dot with a white centre, total 1-degree radius, indicated in **Fig. 1B**) to help the marmosets to associate different stimuli to different tasks.

Trial initiation criteria were similar to the gap task (**Fig. 1B**, top graph). After successful initial fixation, a peripheral target was flashed either to the right or left of the centre fixation stimulus for around 67 ms (**Fig. 1B**, second graph, similar target presentation duration to Hikosaka and Wurtz 1983; Kato et al. 1995; Rivaud-Péchoux et al. 2000; Takikawa et al. 2002; Loe et al. 2012; Willeke et al. 2019; Monosov et al. 2008). The subject had to maintain steady fixation during the delay period (**Fig. 1B**, third graph) until the fixation stimulus offset (go cue). The subject was then required to make a saccade towards the flashed/remembered target location within 1000 ms and maintain gaze for 17 ms to attain a drop of liquid reward along with a rewarding photo that was displayed at the target location. (**Fig. 1B**, bottom graph). The short target fixation time (17 ms) was used to motivate the marmosets. The trial was aborted for similar reasons as in the gap task, and as with the gap task, a one-frame red screen flash was presented as an indication of trial failure. One session was terminated if either 16 successful trials (see Result section) were collected or if a maximum of 32 trials (including error) were collected.

The number of sessions per day was determined based on the marmosets’ level of willingness to work, but in general, an average of 4 sessions of gap and 4 sessions of ODR tasks were performed a day and each session was structured with a different gap/delay period and a different target eccentricity. The liquid reward used for marmosets was prepared by mixing baby supplement banana pudding (Kewpie Corp., Japan) with banana-flavoured Mei-balance (Meiji Holdings Co., Japan).

Human subjects performed the same tasks albeit without the liquid reward. Both tasks were explained to human subjects once and they were told to look at the visual target when it appears in the gap task and the flashed/remembered target location after fixation offset in the ODR task. No further instructions were given to them, in an attempt to make the experimental conditions resemble that of marmosets.

### Saccade Detection

For saccade detection, we used U’n’Eye, a deep neural network developed for eye movement detection and classification (Bellet et al. 2019). The eye data were first pre-processed and the ground-truth saccade dataset was labelled and manually checked following the same procedure and criteria as (Chen et al. 2021). After labelling around 200 trials, the data were gathered and fed into U’n’Eye for network training and validation to get optimal detection. After training, the rest of the pre-processed eye data were fed into the trained network for saccade labelling and the results were inspected by the experimenter to check for mislabelling.

### Data Analysis

#### Definitions of Successful Trials, Error Trials and Saccade Reaction Time

Trials in each task were first separated into either successful or error trials (**Fig. 2)**. A trial was defined as successful if it satisfied the following criteria. First, the saccade that left fixation was generated after the go cue (target onset or fixation offset in gap or ODR tasks, respectively). Second, the target was reached regardless of using one saccade or more. Finally, the trial ended with a reward photo display. On the contrary, a trial was defined as an error if it ended with the one-frame red screen flash presentation.

**Figure 2.**
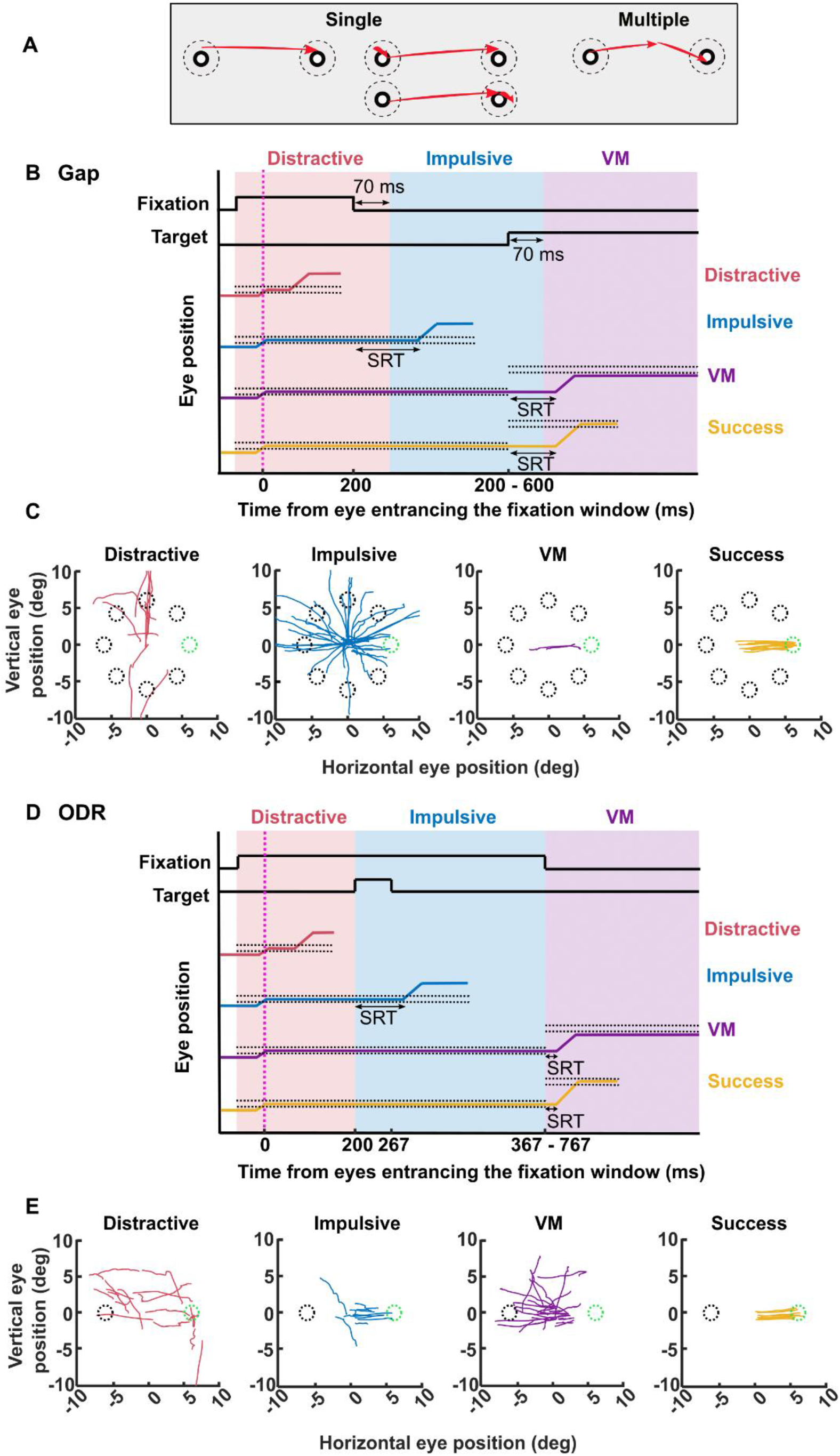
Illustration of task events and the grouping of successful and error trials. **A)** Successful trials were subdivided into a single saccade group (left) or multiple saccades group (right), based on how the subject reached the target stimulus. Dashed circles are the eye windows as described in Fig. 1. Red arrows illustrate saccades made after target onset. **B, D)** Illustration of how we defined successful and error trials in the gap (B) or the ODR (D) tasks based on when the mistake occurred relative to the monitor events. Black solid lines indicate fixation on time and target on time, and coloured lines indicate the relative eye position of the successful and error trial groups (distractive error in red, impulsive error in blue, visuomotor (VM) error in purple, and successful trial in yellow), top to bottom in the figure. Black dashed lines indicate the eye windowing position. The magenta dashed line indicates the time that the subject’s eye position enters the fixation eye window. Illustrations of how we calculated the saccade reaction time (SRT) of the successful and the error trials are also indicated. **C, E)** Example saccade traces extracted from each error group (colour-coded) and successful single saccade group in the gap (C) or the ODR (E) tasks. Saccade traces belong to marmoset S performing the tasks at 6° target eccentricity and 400 ms gap or delay period. Dashed circles indicate potential target locations for each trial. In this example, only the trials where the target was on the right side (indicated as a green dashed circle) were plotted. Refer to **METHOD** for a more detailed explanation.

We further divided successful trials based on how the target was reached: i.e. did the subject use one saccade (single saccade group) or more (multiple saccade group). A trial was defined as a single saccade trial if the saccade started from within the eye window centred around the fixation stimulus (fixation criteria), and landed within the eye window centred at the target location until the reward photo display (target landing criteria), regardless of whether it was escorted with microsaccades or not **(Fig. 2A**, left). Alternatively, a trial was defined as a multiple saccades trial if the first saccade to leave the fixation stimulus meets the fixation criteria and the last saccade to reach the target stimulus meets the target landing criteria **(Fig. 2A**, right). Saccadic reaction time (SRT) was calculated as the time between the target onset in gap task or fixation offset in ODR task and the onset of the saccade (**Fig. 2B, 2D**, yellow eye traces). We used only the single saccade group for SRT and main sequence analysis because it wasn’t clear which saccade was the one intended to answer in the multiple saccades group.

On the other hand, error trials were divided into three sub-groups, distractive error, impulsive error, or visuomotor (VM) error, based on when the trial was terminated relative to the monitor events in each task **(Fig. 2B, 2D)**. For gap task, error trials were labelled based on three monitor events, fixation stimulus onset, fixation stimulus offset, and target stimulus onset. If a trial ended after fixation stimulus onset and before fixation stimulus offset, it was defined as a distractive error trial (**Fig. 2B**, red eye traces). If a trial ended after fixation stimulus offset and before target stimulus onset, it was defined as an impulsive error trial (**Fig. 2B**, blue eye traces). Finally, if a trial ended after the target stimulus onset but with the one-frame red screen flash, it was defined as a VM error trial (**Fig. 2B**, purple eye traces). By definition, impulsive error includes a temporal error (fixation break) and VM error trials includes spatial errors (saccades that ended outside the target eye window) and temporal error (no fixation disengagement, insufficient fixation time on the target and long SRT).

We then looked for the first saccade in the labelled trial that broke fixation (i.e., the saccade started within the fixation window and ended outside the fixation window) **(Fig. 2B)**. This saccade was taken as the saccade that responded to the preceding monitor event in this trial **(Fig. 2C)**, and the SRT was calculated. To be more specific, impulsive error SRT was calculated as the time between fixation offset and the onset of the saccade and VM error SRT was calculated as the time between target onset and the onset of the saccade.

However, anticipatory saccades in successful trials were reported in marmosets performing the gap task (Chen et al. 2021). Because the SRT of the anticipatory saccades was short (< 70 ms), they were likely to be the saccades responding to the fixation stimulus offset (the second nearest monitor event). Considering that, we redefined successful trials and VM error trials with SRT less than 70 ms to be impulsive error trials and impulsive error trials with SRT less than 70 ms to be distractive error trials and recalculated the SRT based on the second nearest monitor events **(Fig. 2B)**. An example of saccades extracted from different error trials and successful trials from marmoset S is shown in **Fig. 2C**.

The definitions of the error trials and SRT calculation for the ODR task were similar to those of gap task **(Fig. 2D)**. The three monitor events were identified as fixation stimulus onset, target stimulus onset, and fixation stimulus offset. Distractive error trials were defined as the trials that ended after fixation stimulus onset and before target stimulus onset (**Fig. 2D**, red eye traces). Impulsive error trials were defined as the trials that ended after target stimulus onset and before fixation stimulus offset (**Fig. 2D**, blue eye traces). VM error trials were defined as the trials that ended after fixation stimulus offset but with the one-frame red screen flash presentation (**Fig. 2D**, purple eye traces). Again, the SRT was calculated for each trial. Impulsive error SRT was calculated as the time between the target onset and the onset of the saccade whereas VM error SRT was calculated as the time between fixation offset and the onset of the saccade. No anticipatory saccades were reported previously, so we did not redefine any trials. An example of saccades extracted from different error trials and successful trials from marmoset S is shown in **Fig. 2E**.

For both tasks, if the subject never entered the fixation eye window after fixation stimulus onset, the trial was defined as a distractive error trial and no saccade or SRT was identified. If the subject did not leave the fixation window after the go cue, the trial was defined as a VM error trial and no saccade or SRT was identified.

The error nomenclature we proposed here reflects our interpretations regarding the possible causes behind each error. For instance, distractive error is related to failure in initiating a trial or breaking fixation at a very early stage of the task, indicating a lack of attention or being distracted enough to not initiate the trial. Impulsive error is related to premature responses (executed before the go cue) and the failure to suppress such responses. Finally, VM error contains incorrect responses, even though the conditions were ripe to produce a correct response (information about the saccadic target was already provided) (Parẻ and Munoz 2001), reflecting either a memory fade (ODR task), or temporal or spatial faults, and hence, a visuomotor related problem.

### Identification of SRT Distribution Components

To identify the SRT components that we observed in the SRT distribution in **Fig. 4**, we first applied data smoothing using the MATLAB function *“smoothdata”*. We used the moving median filter with a sliding window of 2 for humans and 3.9 for marmosets to obtain a smoothed SRT distribution. We then searched for the local minima of the smoothed data using the reversed MATLAB function *“findpeaks”*.

### Identification of Saccades Towards and Away from the Target

To categorize saccades as being directed towards or away from the target, a line through the origin and perpendicular to the extension line connecting the origin and the target coordinates was drawn, dividing the Cartesian coordinate plane into two equal parts. If the saccade endpoint was on the same side as the target coordinates, it was defined as towards the target, whereas if the saccade endpoint was on the side opposite to the target coordinates, it was defined as away from the target.

### Definition of Dysmetric Saccades

VM error saccades directed towards the target were further grouped into hypometric, hypermetric and orthogonal saccades based on the saccades’ endpoints relative to the location of the presented target. To do so, we first identified a deviation vector to have its starting point as the target coordinates and its ending point as the actual saccade endpoint. We also identified a target vector to have its starting point as the origin and its endpoint as the target coordinates. We then calculated the angle between the deviation vector and the target vector. We divided the space around the target into 4 equal quadrants, with quadrant borders oriented 45° from parallel or orthogonal to the origin-target vector (**Fig. 6**). A saccade was defined as hypometric (under) if it fell within the quadrant on the near side of the target, and hypermetric (over) if it fell within the quadrant on the far side of the target. Finally, a saccade was defined as orthogonal if the angle was in either of the other two quadrants.

### Saccade Main Sequence

We analysed the saccade main sequence to characterize the changes in saccade kinematics for different trial groups (Bahill et al. 1975). We first plotted the saccade amplitude (x) against the saccade peak velocity (f(x)). We then fitted the data with the following equation:

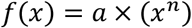

Where a and n are free parameters representing the slope and the exponent.

### Statistical Analysis

All the statistical analyses were performed after we grouped the collected data according to gap/delay period as short (< 150 ms), intermediate (≥ 150 and < 300 ms) and long (≥ 300 ms) for each eccentricity used (4,6 or 8 deg in gap task and 4 or 6 deg in ODR task). The statistical tests and results are described in the main text and the figure legends.

## RESULTS

### Marmosets were More Prone to Making Mistakes than Humans

We first collected eye movement data from 3 humans and 3 marmosets performing the gap and ODR tasks in parallel (**Fig. 1**, averaged trials per marmoset: 3368 in gap task, 6649 in ODR task; per human: 4237 in gap task, 2650 in ODR task). These trials were separated into 2 types of successful trials (Fig. 2A) and 3 types of error trials **(Fig. 2B - 2D)** according to the criteria defined in **METHOD**. We compared between gap and ODR tasks performance and how often each successful and error trials (**METHOD, Fig. 2B, 2D**) occurred in marmosets and humans.

In marmosets, as indicated in **Fig. 3A, 3B**, the overall error trial ratio averaged across marmosets was higher in ODR task (77.7%) than in gap task (21%).

**Figure 3.**
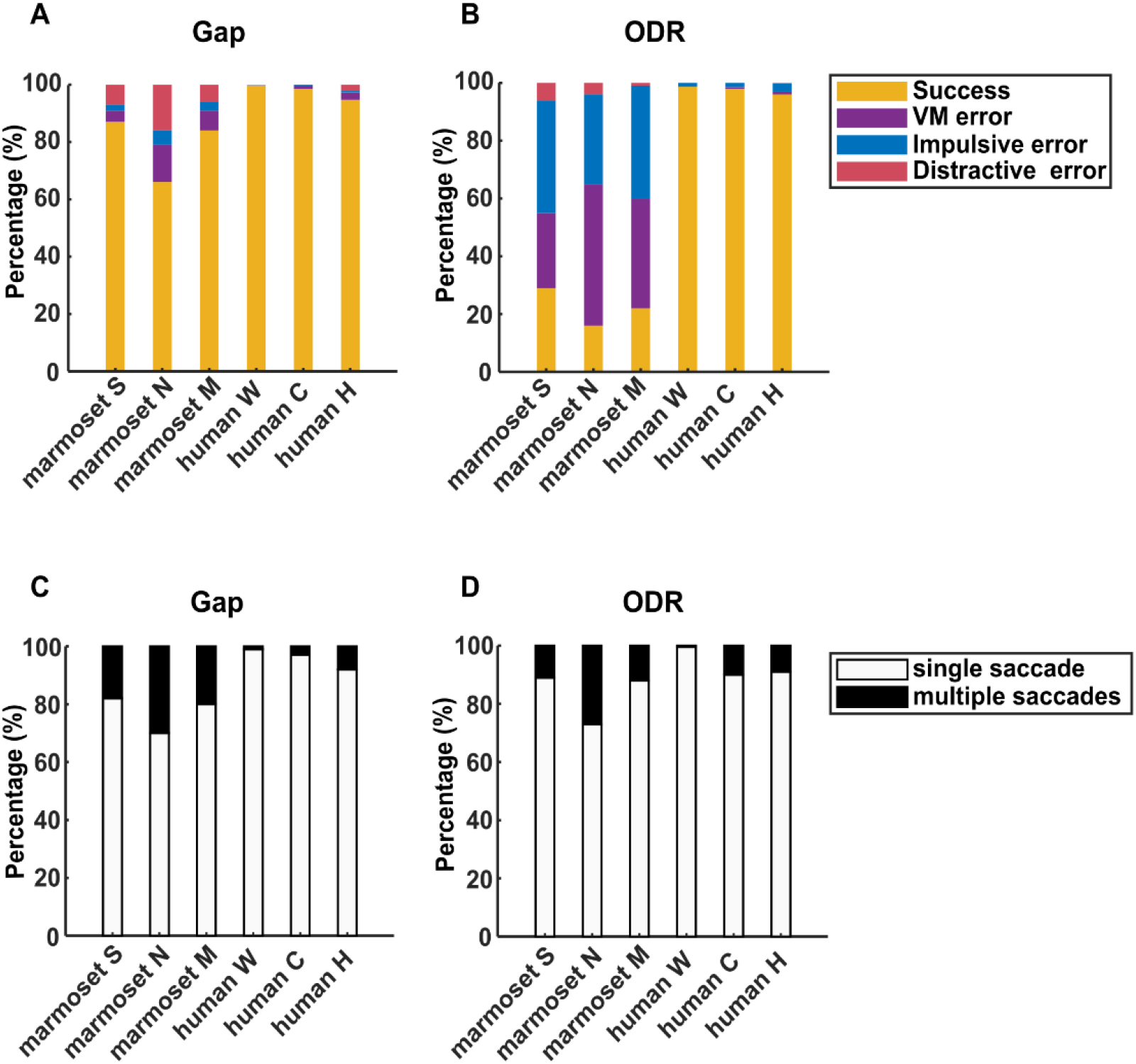
Percentage of each subgroup of successful and error trials in gap and ODR tasks. Trials collected from all subjects are shown here. **A, B)** Success and error ratios are shown in (A) for the gap task and (B) for the ODR task. Comparing gap and ODR tasks, marmosets’ data shows that the distractive error ratio was significantly higher in gap task (p=0.01). Impulsive and visuomotor (VM) errors were significantly higher in ODR task (p=2.86×10^−8^, 2.5×10^−8^ respectively), whereas the success ratio was significantly higher in gap task (p=1.9×10^−8^). Humans made very scarce errors and therefore, no statistical analysis was performed, but humans’ success ratio in both gap and ODR tasks significantly outweighed that of marmosets (p=9.98×10^−10^, p=3.2×10^−7^ respectively). **C, D)** Subgroups of successful trials are shown for gap (C) and ODR (D) tasks. Single saccade or multiple saccades groups were not significantly different between the tasks in marmosets (p=0.22, 0.22 respectively) or humans (p=0.38, 0.94 respectively). Wilcoxon rank-sum test was used for statistical analyses.

**Figure 4.**
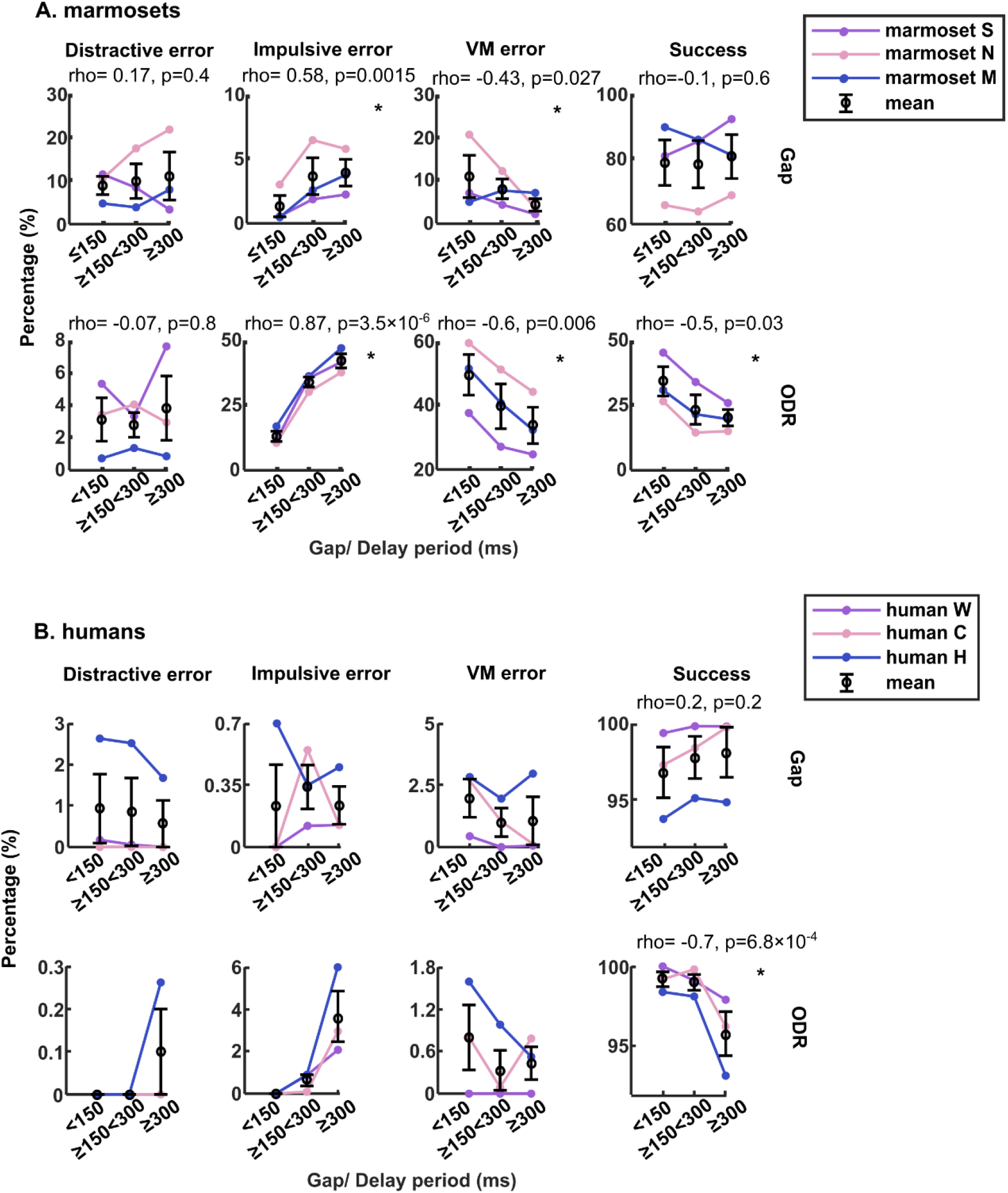
Change in success and error ratios with gap or delay periods in marmosets and humans (B) in both tasks. **A)** The trials proportions for impulsive and visuomotor (VM) error and successful trials against short (<150 ms), medium (150-250 ms), and long (> 250 ms) gap/delay periods were plotted for marmosets separating gap (top row) or ODR task (bottom row) to see the correlation. Statistics are provided in each subfigure and the ones with significant correlation are marked with *. **B)** The same analysis was also performed on humans’ data. We didn’t perform statistical tests on human errors due to the scarcity of error trials. But please note that a similar trend of increasing impulsive error and decreasing VM error with longer gap/delay periods was observed in both marmosets and humans. We found no correlation between the success ratio and gap period, whereas a negative correlation with the delay period was statistically confirmed in both species. Spearman linear correlation test was used for statistical analysis.

We further compared the frequency of each error type in both tasks. We expected to see a higher ratio of distractive error in ODR task as the task is more demanding and might induce more opt-outs than in gap task. However, the distractive error ratio was lower in all three marmosets in ODR task (on average 3.6% in ODR task vs 9.7% in gap task, p=0.01, Wilcoxon rank-sum test, **Fig. 3A, 3B)**.

As for the impulsive error, we hypothesized that the marmosets would be more susceptible to impulsive responses in ODR task because the target was briefly displayed before the delay period in ODR task, whereas it was not yet shown during the gap period in gap task. **(Fig. 2C, 2E)**. Indeed, we found that the impulsive error ratio was higher in ODR task (36.3%) than in gap task (3.3%, p= 2.86×10^−8^, Wilcoxon rank-sum test, **Fig. 3A, 3B**). We also expected a higher VM error ratio in ODR task due to the absence of the visual target, and the results met our expectations (37.7% in ODR task vs 8% in gap task, 2.5×10^−8^, Wilcoxon rank-sum test, **Fig. 3A, 3B**).

Importantly, even though humans made few errors in both tasks (2.3% and 2.4% in gap and ODR tasks respectively) and we were unable to perform statistical comparisons between the tasks, a lower distractive error ratio (0.03% in ODR task vs 0.7% in gap task) and a higher impulsive error ratio (1.9% in ODR task vs 0.4% in gap task) in ODR task were observed, similar to the marmosets.

On the other hand, we analysed how often the successful trials were answered with a single vs multiple saccades (**METHOD, Fig.2A**). We found that the marmosets showed a higher probability to reach the target stimulus with a single saccade in both tasks (gap: 77.3%, ODR: 83.3%). The likelihood of answering with either a single saccade or multiple saccades was not significantly different between the two tasks (gap: p=0.2, ODR: p=0.2, Wilcoxon rank-sum test, **Fig. 3C, 3D)**. Humans showed similar results in both tasks as well (gap: 96%, p=0.38, ODR: 93.5%, p=0.94, Wilcoxon rank-sum test).

### Marmosets Learned the ODR Task

Before proceeding further with dissecting the error characteristics, we were concerned that the marmosets’ overall performance in ODR task was not as good as in gap task. We wondered if this was just a consequence of the higher task difficulty, or because the marmosets didn’t comprehend the task requirements (i.e. when and where to make a response).

To first answer if the marmosets understood when to make a response, we plotted the relative frequency histogram and the cumulative probability distribution of the SRT for all the saccades made after target onset (i.e. saccades collected from impulsive error, VM error and successful trials) from marmoset S as an example **(Supplementary xsFig. 1A)** and human H **(Supplementary Fig. 1B)** in the same manner as a reference.

By analysing the peaks and troughs of the relative frequency histogram of the SRT distribution, we could identify a maximum of four distribution components (labelled1-4 in **Supplementary Fig. 1A, Supplementary Fig. 1B**, top-left illustration), two before and two after the fixation stimulus offset (indicated as the magenta line). These components were further confirmed by applying trough detection (**METHOD, Supplementary Fig. 1**, orange curves and blue stars).

The most important components regarding our question are the ones around the fixation offset, components 2 and 3, and they exhibited a strong discontinuity in the SRT histogram and cumulative distributions. Interestingly, the SRT distribution and discontinuity after target onset from the marmosets captured most of the characteristics of the human subjects This discontinuity of the SRT distribution and the cumulative distribution around the fixation offset indicates that saccades before and after the magenta line are likely responding to different monitor events, signifying that the marmosets understood when to make a response.

We next examined whether the marmosets understood where to respond rather than just gazing randomly or guessing either left or right. To do so, we examined the VM error and the successful saccades’ landing positions (the saccade that reached the target) for each marmoset **(Supplementary Fig. 2)**. We mirrored the left targets and their corresponding saccades landing positions to the right side. The inside of the right solid circle in the figure represents the correct response whereas the left dashed circle represents the incorrect response. If the marmosets were guessing where was the target, a similar amount of saccades’ landing positions should show up in both the solid and the dashed circles. However, as the figure shows, a significant majority of the saccades landed inside the right circle, (4° target: p= 1.1×10^−9^, 9.2×10^−4^ and 4.2×10^−9^, for marmoset S, N and M ; 6° target: p=3.9×10^−50^, 8.3×10^−20^, and p= 2.1×10^−35^, for marmoset S, N and M, Wilcoxon rank-sum test) signifying that the marmosets’ successful or unsuccessful saccades were not guessing but rather directed according to the task instructions.

The above results indicate that the lower performance in ODR task was mainly due to the task difficulty and not because the marmosets didn’t comprehend the task structure.

### Longer Gap/Delay Periods Correlated with Higher Impulsive and Lower VM Error Frequencies

Based on the previous results that confirmed marmosets’ comprehension of ODR task, we were confident to proceed and investigate how the varied task conditions (gap/delayed period and target eccentricities) dictated the performance in both species **(Fig. 4)**.

The first condition that varied from session to session was the gap/delay period in gap/ODR tasks. By analysing the correlation between gap/delay period and the ratio of errors or successful trials, we expected to see a positive correlation between gap/delay and impulsive error ratio because chances to break fixation increase with a longer waiting time. Indeed, we found that the impulsive error ratio increased with longer gap/delay periods in both tasks (gap: p=0.0015, r=0.58, ODR: p=3.5×10^−6^, r= 0.87, Spearman linear correlation) (**Fig. 4A**, second column). Interestingly, the VM error ratio showed a negative correlation with a longer gap/delay period (gap: p=0.027, r= -0.43, ODR: p=0.006, r= -0.6, Spearman linear correlation) (**Fig. 4A**, third column). Furthermore, we found that the success ratio didn’t show consistent change with the gap period (p=0.6, Spearman linear correlation), however, it showed a significant negative correlation with the delay period in ODR task (p=0.03, r= -0.5, Spearman linear correlation) **(Fig. 4A**, last column).

Although the error trials were too less to perform statistical tests for humans, we observed a similar pattern to what we observed in marmosets (**Fig. 4B**). In other words, increasing impulsive error ratio and decreasing VM error ratio with a longer delay period in ODR, decreasing success ratio with a longer delay period in ODR task (p=6.8×10^−4^, r= -0.7).

To analyse the correlation between marmosets and humans, we performed a linear correlation test between the two species and found that both species showed a significant correlation in impulsive and VM error trends in ODR task (r= 0.75, p= 0.018; r=0.81, p= 0.009 respectively), and success in gap task (r= 0.8, p= 0.005, **Table 1**).

**Table 1.** Linear correlation test results. The correlation (r) and the p-value (p) of the linear correlation test between marmosets and humans.

| Success and Error percentage vs gap/delay period (related to Fig. 4) |  |  |  |  |  |
| --- | --- | --- | --- | --- | --- |
| Gap task |  |  | ODR task |  |  |
| Impulsive error | VM error | Success | Impulsive error | VM error | Success |
| r= 0.4 p= 0.29 | r= 0.43 p= 0.25 | r= 0.8 p= 0.005 | r= 0.76 p= 0.018 | r= 0.81 p= 0.009 | r= 0.6 p= 0.07 |
| Success and Error percentage vs target eccentricity (related to Supplementary Fig. 4) |  |  |  |  |  |
| Gap task |  |  | ODR task |  |  |
| Impulsive error | VM error | Success | Impulsive error | VM error | Success |
| r= 0.5 p= 0.16 | r= 0.46 p= 0.21 | r= 0.9 p= 9.8x10<sup>-4</sup> | r= 0.7 p= 0.11 | r= 0.47 p= 0.34 | r= 0.55 p= 0.26 |
| SRT vs gap/delay period (related to Fig. 5) |  |  |  |  |  |
| Gap task |  |  | ODR task |  |  |
| Impulsive error | VM error | Success | Impulsive error | VM error | Success |
| r= 0.98 p= 1.45x10<sup>-6</sup> | r= 0.61 p= 0.08 | r= 0.86 p= 0.0028 | r= 0.23 p= 0.56 | r= 0.54 p= 0.13 | r= 0.7 p= 0.04 |
| SRT vs target eccentricity(related to Supplementary Fig. 5) |  |  |  |  |  |
| Gap task |  |  | ODR task |  |  |
| Impulsive error | VM error | Success | Impulsive error | VM error | Success |
| r= 0.5 p= 0.18 | r= 0.2 p= 0.58 | r= 0.3 p= 0.4 | r= 0.8 p= 0.045 | r= 0.7 p= 0.11 | r= 0.88 p= 0.02 |

The second condition that varied from session to session was the target eccentricities. In marmosets, we didn’t see much influence of the target eccentricity on either success or failure except for a positive correlation with the VM error ratio in gap task (p= 0.04, r= 0.4, Spearman linear correlation) and a negative correlation in ODR task (p= 0.019, Wilcoxon signed-rank) **(Supplementary Fig. 3A**, third column). In humans, a similar trend of increasing VM error ratio with farther eccentricities in gap task was observed (**Fig. 4B, Supplementary Fig. 3B**, third column). Linear correlation test between the species showed a significant correlation between trends observed in success in gap task (r=0.9, p= 9.8×10^−4^, **Table 1**).

### Longer Gap/Delay Periods Correlated with a Longer Impulsive Error SRT and a Shorter Success SRT

Next, we examined how the varied task conditions altered the SRT. Because SRT reflects the time needed for the sensory-to-motor process to occur, we wanted to investigate if such a process differs when mistakes were made or when correct vs incorrect responses were made **(Fig. 5)**.

**Figure 5.**
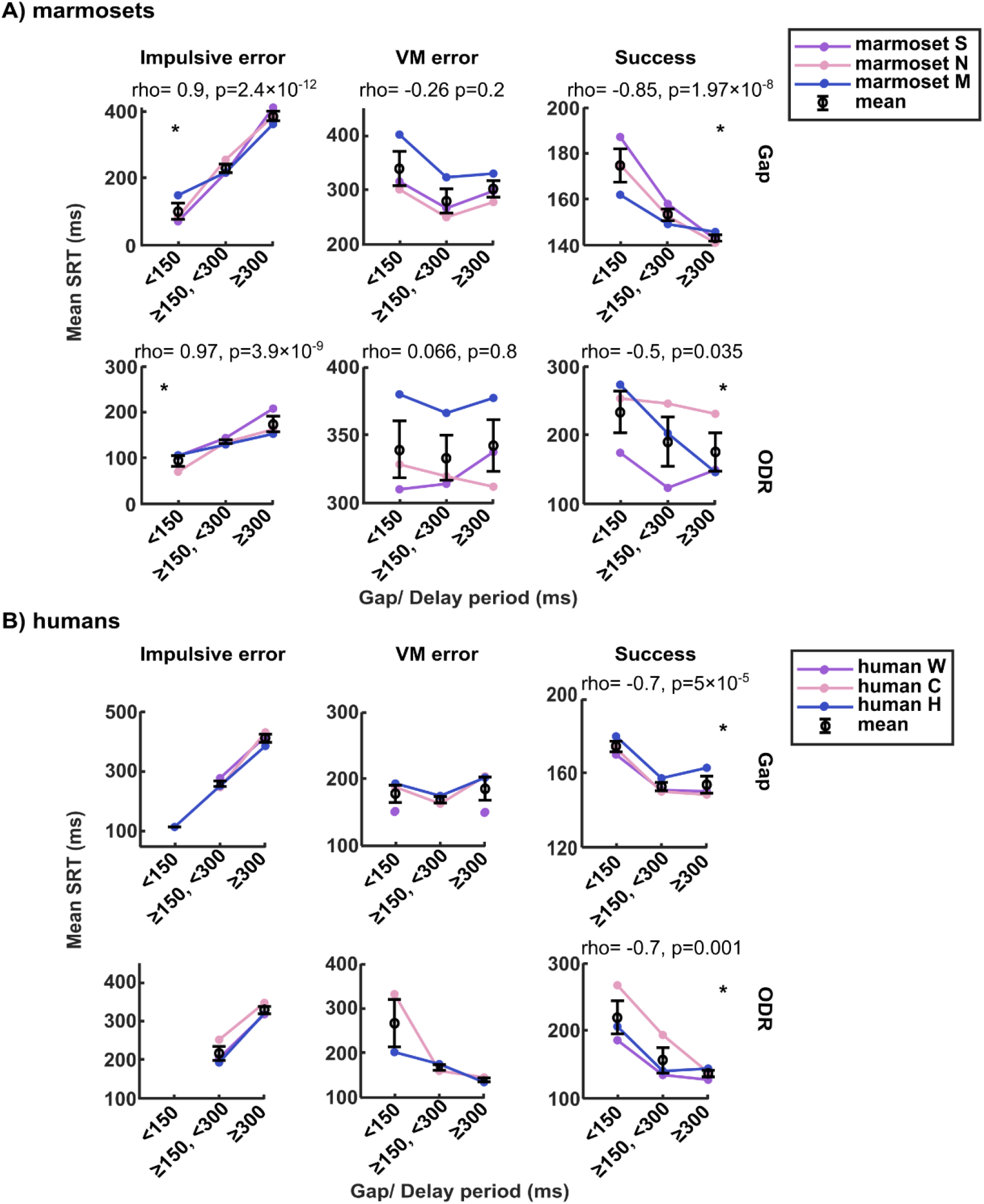
Change in saccade reaction time (SRT) of success and error subgroups with gap or delay periods in marmosets and humans in both tasks. **A)** SRT of impulsive and visuomotor (VM) error and successful trials against short (<150 ms), medium (150-250 ms), and long (> 250 ms) gap/delay periods were plotted for marmosets, separating gap (top row) and ODR (bottom row) tasks to see the correlation. The Spearman linear correlation test results are provided in each subfigure and the ones with significant correlation are marked with *. We also compared the SRTs among success and error subgroups. SRT of impulsive error in the ODR task was shorter than in gap task (p= 0.002, Wilcoxon rank sum test). **B)** The same analysis was also performed on humans’ data. We didn’t perform correlation statistical tests on human errors due to the scarcity of error trials. But please note that a similar trend of increasing impulsive error SRTs with longer gap/delay periods was observed in both marmosets and humans. Decreasing successful SRTs with longer gap/delay periods in both tasks was statistically confirmed in both species.

For both marmosets and humans, impulsive error SRT was, in general, shorter in ODR task (marmoset: 100-200 ms; human: 200-300 ms) than in gap task (marmoset: 100-400 ms; human: 300-400 ms; **Fig. 5A, 5B**, left column). As described previously in **Supplementary Fig. 1** for ODR task, this was due to the first two components in the SRT distribution, with component 1 being triggered by the flashed target and component 2 by the urgency to make a response. This could also explain why the saccades were mainly directed towards the target (**Fig. 6A**). However, in gap task, all the impulsive error saccades were anticipatory, which natively happened after the peripheral target onset. Due to the above reasons, the impulsive error SRT of either gap or ODR task should have naturally a positive correlation with gap or delay period, respectively, (gap: r: 0.9, p= 2.4×10^−12^, ODR: r: 0.97, p=3.9×10^−9^, Spearman linear correlation). These results indicate that the presence of a visually accessible target facilitates fixation disengagement and the urgency to make a response. We also noticed a significant negative correlation with successful SRT in both tasks and both species **(Fig. 5A, 5B**, right column), (marmosets; gap: r: -0.85, p= 1.97×10^−8^, ODR: r: -0.5, p= 0.035; humans; gap: r: - 0.7, p= 5×10^−5^, ODR: r: -0.7, p= 0.001). In addition, the linear correlation test between the species showed a significant correlation between the trends observed in success in both tasks (gap: r=0.86, p=0.003; ODR: r=0.7, p=0.04) and impulsive error in gap task (r= 0.98, p= 1.45×10^−6^, **Table 1**).

**Figure 6.**
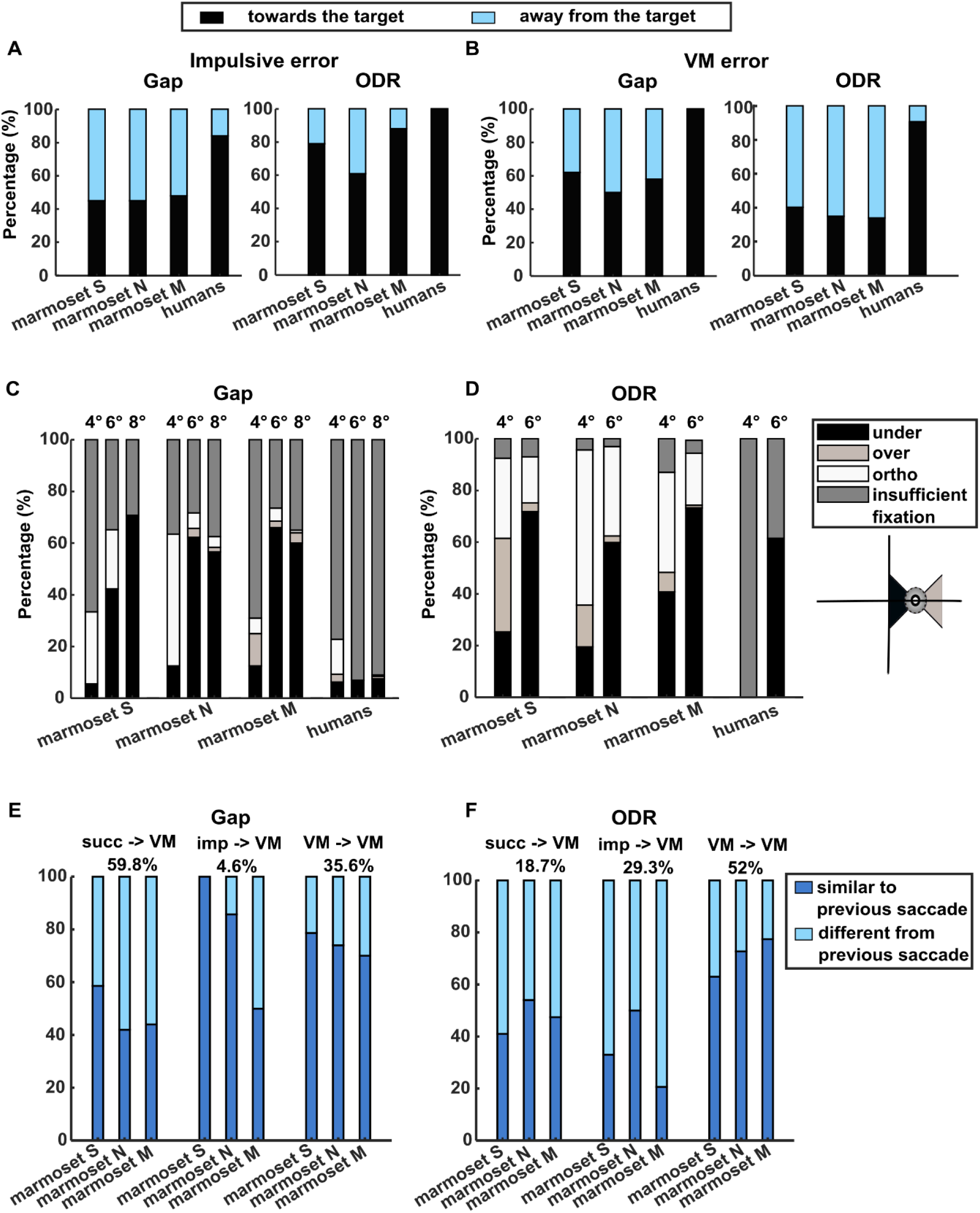
Grouping of error trials according to the saccade direction relative to the target location or the previous trial decision. Saccades from the impulsive and visuomotor (VM) error trials were plotted and analysed here for both marmosets and humans. Although we plotted the figure with all humans’ results combined, the statistical tests were performed for them separately, similar to marmosets. **A)** In marmosets, saccades from impulsive error trials were separated based on going toward or away from the target and the marmosets showed an equal proportion of saccades in gap task but more saccades going toward the target in ODR task (p=0.2, p=7.3×10^−7^, respectively, Wilcoxon rank sum test). However, humans showed more saccades going towards the target in both gap and ODR tasks (p=1.6×10^−6^, p=5.5×10^−6^, respectively, Wilcoxon rank sum test). **B)** The same analysis on VM error trials in marmosets showed an equal proportion of saccades in the gap but more saccades going away from the target in the ODR task (p=0.4, p= p=3.2×10^−7^, respectively, Wilcoxon rank sum test), whereas again humans show more saccades going towards the target in both gap and ODR tasks (p=1.3×10^−9^, p=5.1×10^−5^, respectively, Wilcoxon rank sum test). **C, D)** Further separation of saccades directed towards the target in the VM error trials into undershooting (under), overshooting (over), or orthogonal (ortho), or saccades landed within the eye window but with insufficient fixation. A positive correlation between the target eccentricity and saccade undershooting was observed in both gap and ODR tasks (r=0.7, p=2.2×10^−5^, Spearman linear correlation test; p=0.004, Wilcoxon sign rank test, respectively) in marmosets. However, the correlation between the target eccentricity and saccade overshooting was only observed in ODR task (gap: r=0.2, p=0.36, Spearman linear correlation test; ODR: p=0.008, Wilcoxon sign rank test, respectively). **E, F)** Further separation of saccades directed away from the target in the VM error trials based on the direction of the previous saccade. The total trial percentage combining the three marmosets is shown below each condition. If the VM trial was preceded by a VM experience, marmosets significantly repeated the same answer in ODR task (gap: p=0.02, ODR: p= 3.2×10^−7^, Wilcoxon rank sum test). If the VM trial was preceded by an impulsive error experience, the saccade from the previous trial was directed in the opposite direction of the current saccade in ODR task (p=1.9×10^−4^, Wilcoxon rank sum test), but in the same direction in gap task (p=0.04, Wilcoxon rank sum test). Finally, if the VM trial was preceded by a successful experience, the previous and the next saccades were directed in the same or opposite direction of each other with almost equal probability, in both gap and ODR tasks (p=0.43, p=0.96, respectively, Wilcoxon rank sum test).

In addition, we provide the boxplot figure of the SRT **(Supplementary Fig. 4)** because most of the SRTs were not normally distributed (**Supplementary Fig. 1**). Similar to what was shown in **Supplementary Fig. 1**, we observed that both marmosets and humans showed a very short SRT (immediately following the fixation offset) in both the VM error and the successful trials in ODR task, indicating that the fixation stimulus offset was predicted in both groups of trials (**Supplementary Fig. 4**). We also analysed the correlation using the median of the SRTs and the results were similar to that using the mean.

On the other hand, the target eccentricity only significantly influenced marmosets’ successful SRT in ODR task only (marmosets: p= 0.004, humans: p=0.098, Wilcoxon signed-rank test) (**Supplementary Fig. 5**). Linear correlation test between the species showed a significant correlation between the trends observed in success in ODR task (r=0.88, p=0.02), (**Table 1**).

### Target Visibility Induced Impulsive Error Saccades in ODR Task

Although the positive correlation between the gap/delay period and the impulsive error frequency met our expectations, the negative correlation with the VM error frequency suggests mechanisms other than merely not detecting the target in gap task or having memory fade out in ODR task **(Fig. 4, Supplementary Fig. 4)**, as both suggest opposite outcomes. To further investigate the underlying reasons, we separated those error groups depending on whether the saccade vector was directed towards or away from the target.

We found that across all three marmosets, impulsive error saccades made in gap task were directed 46% toward and 54% away from the target stimulus, with almost equal probability (p=0.2, Wilcoxon signed-rank test) **(METHOD, Fig. 2C, Fig. 6A)**. We further confirmed that the saccades from the impulsive error trials were all anticipatory saccades (SRT <70 ms) (Becker W. 1989; Dorris and Munoz 1998; Walker et al. 2000). This is similar to a previous report that noted the anticipatory saccades with correct direction and direction errors would be equally frequent (Kalesnykas and Hallett 1987). Although saccades from the impulsive error trials in humans were also anticipatory saccades, they showed a higher tendency of going toward the target (84%, p=1.6×10^−6^, Wilcoxon rank sum test). This result might be affected by the scarcity of the impulsive error trials (34 trials combining all three humans, vs 338 trials combining all three marmosets).

On the other hand, impulsive error saccades made in ODR task showed a significant bias to go toward the target stimulus (76% towards, p=7.3×10^−7^, Wilcoxon rank sum test, **Fig. 6A**). This is likely because the target was briefly presented before the delay **(Fig. 1B)** and the marmosets were incapable of suppressing their reflexive saccades **(Supplementary Fig. 1A)**. Humans showed similar results (100% towards, p=5.5×10^−6^, Wilcoxon rank sum test), yet again, the number of impulsive error trials were scarce (148 trials combining all three humans).

As for the VM error, the marmosets showed an almost equal probability of making a saccade towards (57%) or away (43%) from the target stimulus in gap task, whereas a significant proportion of the saccades were directed away (64%) from the briefly presented target in ODR task (gap: p=0.4, ODR: p= p=3.2×10^−7^, Wilcoxon rank sum test) **(Fig. 6B)**. Humans showed a higher tendency to make saccades directed towards the target in gap task (100%) and ODR task (92.5%), (gap: p=1.3×10^−9^, ODR: p=5.1×10^−5^, Wilcoxon rank sum test). This might designate an interspecies variance, but the human error trials count was very low to be ascertained, (157 trials in gap task, 36 trials in ODR task for humans, vs 870 trials in gap task, 7067 trials in ODR task trial for marmosets).

### Saccade Undershooting Correlated with Target Eccentricity

To further understand why incorrect answers occurred despite of the appearance of target information (VM error), we first examined the landing position and the target fixation time of the VM error saccades directed towards different target eccentricities. Results showed that failure was attributed to dysmetria (overshooting, undershooting, orthogonal saccades) or insufficient fixation on the target (less than 200 ms in gap task or 17 ms in ODR task, **Fig. 6C, 6D**). Even though human subjects didn’t have many errors, among the collected ones, insufficient fixation was the main failure reason in both tasks **(Fig. 6C, 6D)**.

We also found that farther eccentricities were significantly correlated with more saccade undershooting in both tasks in marmosets (gap: p=2.2×10^−5^, r=0.7, Spearman linear correlation test; ODR: p=0.004, Wilcoxon sign rank test). The same analysis in humans showed a similar trend, albeit without sufficient trials for statistical tests. Furthermore, nearer eccentricities correlated with more overshooting only in ODR task in marmosets (gap: r=0.2, Spearman linear correlation test; ODR: p=0.008, Wilcoxon sign rank test).

### Marmosets Repeated the Same Incorrect Answer

The VM error saccades directed away from the target stimulus in marmosets were particularly puzzling to us **(Fig. 2B, 2E, 6B)**. In gap task, the target was already visible by the time a VM error saccade was made. In ODR task, if they forgot the target location, an equal bias of saccading towards or away from the target stimulus would’ve been seen, if they roughly remembered the target location, the saccade direction should’ve been made towards the target. Then, how did this happen? We hypothesized that this could be related to the combination of our task design and the marmosets’ answering strategy. To prevent the marmosets from favouring one spatial location over the others, if a trial was unsuccessful, the following trial will reuse the same target location. However, if the marmosets did not notice this implementation and decided to saccade in the opposite direction because they were not rewarded in the previous trial, a saccade away from the target may occur. Therefore, we analysed the task history by inspecting the saccade direction that preceded the VM error saccade directed away from the target (i.e. consecutive trials only) to see if the previous experience affected their responses. (**Fig. 6E, 6F**).

We found that if the previous trial was successful, saccades were directed either in the same or opposite direction with equal probability in both tasks (48.5%, p=0.43 and 47.5%, p=0.96 same direction in gap and ODR tasks respectively, Wilcoxon sign rank test), meaning that they didn’t favour the direction that delivered the reward. However, if the previous trial was an impulsive error trial, more consecutive saccades had a similar direction in gap task (78.5% same direction, p=0.04), whereas more had a different direction in ODR task (65.5% opposite direction, p=1.9×10^−4^, Wilcoxon sign rank test). Finally, if the previous trial was a VM error trial, marmosets’ decision was significantly biased towards repeating the same answer (away from the target), in both tasks (74.2%, p=0.02 and 71%, p=3.2×10^−7^ same direction in gap and ODR tasks respectively, Wilcoxon sign rank test). Surprisingly, instead of alternating their answer, the marmosets tended to repeat the same answer (saccade direction) in consecutive error trials.

### Distinct Saccade Main Sequence Between Success and Errors Observed

Finally, we analysed the saccade main sequence. Because saccade peak velocity and amplitude are known to be governed by a stereotyped relationship within subjects, we wondered if such a relationship is also preserved among successful and unsuccessful defined saccades **(Fig. 7A, 7B)**. We found that in the gap task, marmosets showed a higher slope and a lower exponent of the fitted amplitude-peak velocity curve parameters for successful saccades compared to unsuccessful ones (**METHOD, Fig. 7C**). A similar result can be observed in humans, although the impulsive error trials were too less causing the confidence intervals to be large **(Table 2)**. On the other hand, impulsive error saccades in ODR task had the highest fitted slope in marmosets, but the difference from other trial types is less obvious. The parameters in humans were not different from each other **(Fig. 7D)**. The confidence intervals were large and overlapped due to the scarcity of error trials in humans.

**Figure 7.**
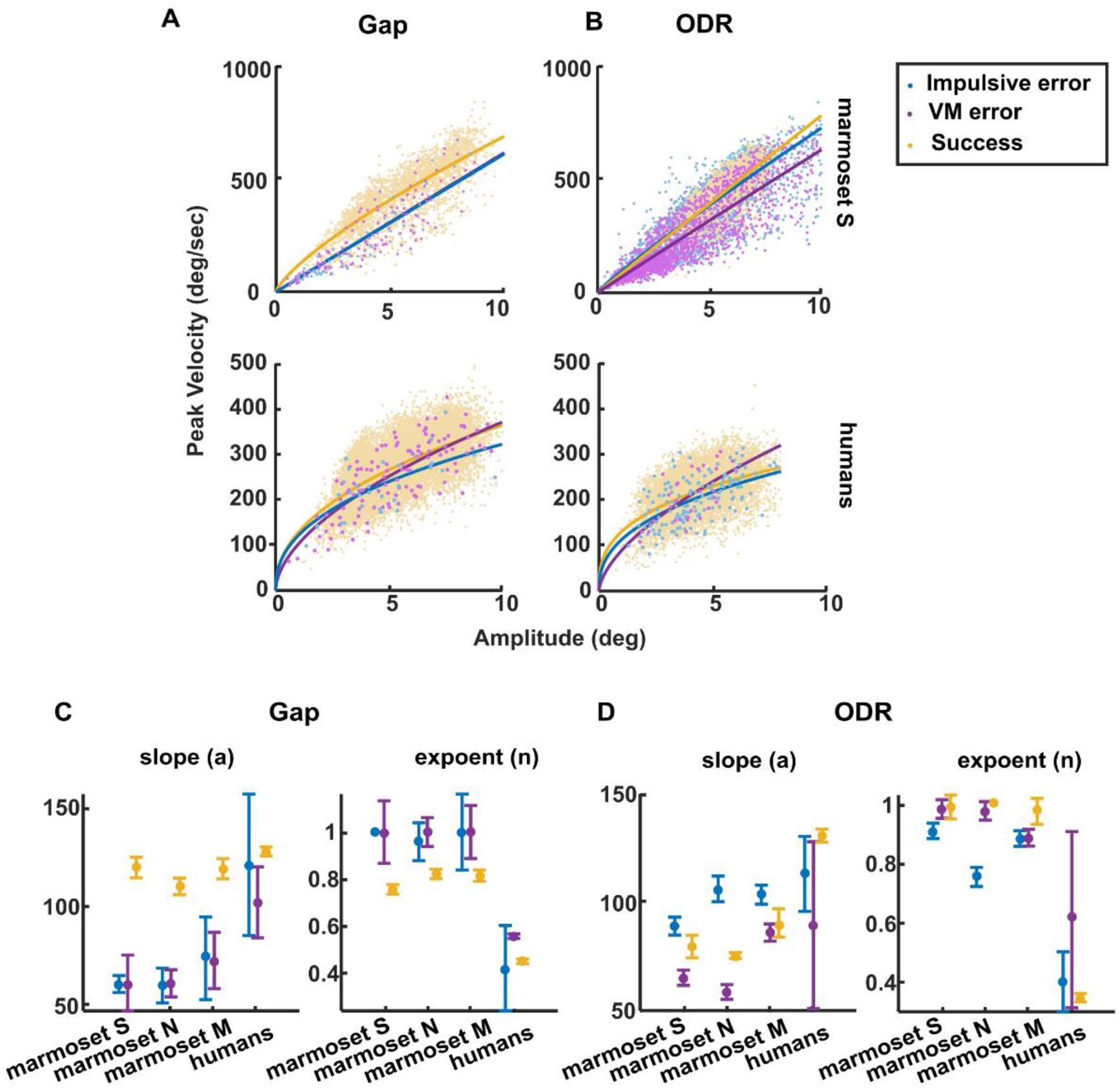
Saccade main sequence of the success and error subgroups in marmosets and humans. **A)** Saccade peak velocities in the gap task were plotted against saccade amplitudes and colour-coded, separating impulsive error, visuomotor (VM) error and successful trials. Marmoset’s S (top) data was plotted as an example, and humans’ data combined (bottom) due to the insufficient number of error trials. Saccade main sequences were fitted using the following fitting equation *f(x)=a*∗*(x*^*n*^). **B)** The same analysis was performed also for the ODR task. **C, D)** The fitting parameters (*a* and *n*, indicating the slope and the curvature, respectively) for each marmoset and combined humans’ data were plotted for gap and ODR tasks. Error bars represent 95% of the confidence interval of the fitted parameter.

**Table 2.**
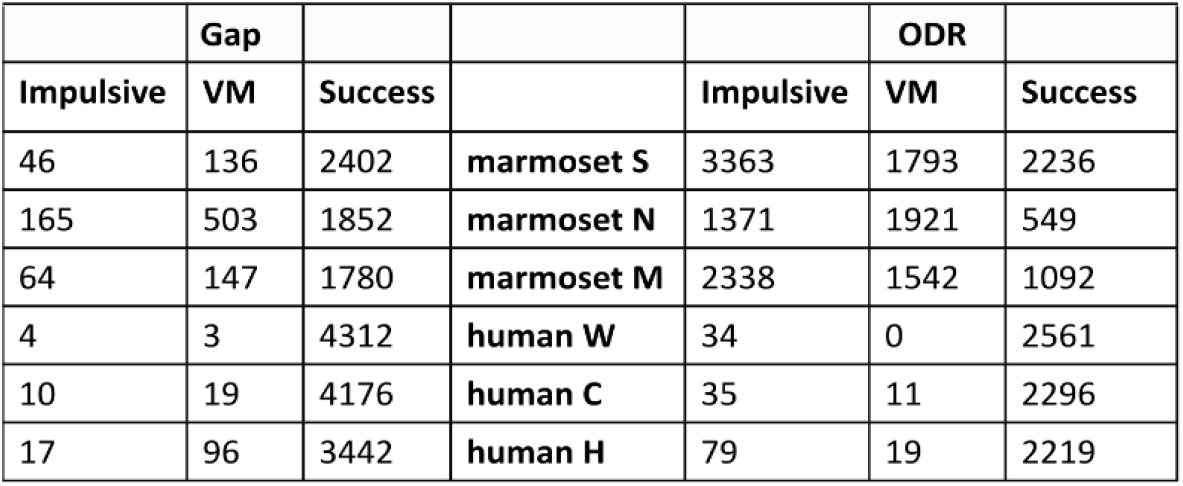
The number of successful and error trials for each subject in both tasks. The number of trials in success and error categories used in saccade main sequence calculations is provided for each subject.

## DISCUSSION

### Dissecting Errors Revealed Similarities Between Marmosets and Humans

In this study, we showed that the marmosets were capable of performing a gap saccade task (in short, gap task) and an oculomotor delayed response (ODR) task. We collected both successful and error trials, categorized error trials into three groups based on the time point the error occurred during the tasks, and analysed them along with successful trials to interpret why subjects made such mistakes. We compared trends between marmosets and humans. We started by looking at the overall performance in each task and found that marmosets and humans made more errors in ODR task than in gap task (**Fig. 3A**), consistent with a previous report on humans (Brown et al. 2004).

Although humans’ error trials were too sparse to perform statistical analyses, we found that marmosets and humans had fewer distractive errors and more impulsive and visuomotor (VM) errors in ODR task compared to gap task. We also observed a similar correlation between the gap/delay period and the error frequencies as well as saccade reaction time (SRT) in both species. The high error ratio in marmosets could be partially explained by their tendency to reiterate the same saccade away from the target. Finally, we found that the saccade main sequence differed between error and success in both tasks

In what follows, we discuss in detail the potential causes of the three error groups (distractive, impulsive, VM errors) based on our analysis in the result section and the new findings in successful trials.

### Distractive Error

We found that the distractive error ratio was higher in gap task in contrast to ODR task in both species (**Fig. 3A**). Despite the scarcity of other direct reports, similar results could be obtained by calculating the breakdown of errors from (Brown et al. 2004). We refer to this more frequent distractive error in gap task as a potential “mind-wandering” state, where the attention is generally directed to task-irrelevant internal thoughts for easy and repetitive tasks (Reichle et al. 2010), which is related to the activation of the default network (medial prefrontal cortex, cingulate cortex, precuneus and angular gyrus) (Mason et al. 2007). When the internal thoughts are activated, the eyes are disengaged from the external stimulus to guard the internal thoughts against external events (Smallwood et al. 2007). Thus, to successfully execute a demanding task, one must stay concentrated on the ongoing task and not go into a mind-wandering state (Benedek et al. 2017). Therefore, the lower distractive error in ODR task might be due to the higher task demand, which activates more cognitive processes. Alternatively, the gap task is mainly driven by external stimuli and could lead to a mind-wandering state easily. Such differences will depend on the type of task rather than its particular parameters, and this is likely why we did not observe a correlation between parameters like gap/delay period or target eccentricities with distractive error rate in either marmosets or humans (**Fig. 4, Supplementary Fig. 3**), which agrees with the findings in after-task phases saccades (i.e. distractive saccades) observed in humans (Takemoto et al. 2022).

On the other hand, we know that the frontal eye field (FEF) is active during visually-guided and memory-guided saccade tasks (Bruce and Goldberg 1985) but the neural discharge differences between planned versus distractive saccades are less clear. However, a recent report of a study on macaques showed that non-goal-directed saccades had a lower firing rate compared to goal-directed saccades, and this appears to be due to the lack of attention while performing the task (Sendhilnathan et al. 2021). These results suggest that FEF neurons responses are lower for unattended targets. Since those non-goal-directed saccades which are induced by looking at visual stimuli without engaging in a specific task bear similarity to our distractive error saccades, we think that FEF is likely less involved during mind-wandering. In addition, superior colliculus (SC) is another essential structure to be considered as it is known to be involved not only in visually driven saccades but also in saccades generated in complete darkness where no FEF firing is preceded (Bizzi 1968). Neural recording data in marmosets would further enhance our understanding of these particular saccades and if they are driven by top-down vs bottom-up processes.

### Impulsive Error

We found that the impulsive error ratio was higher in ODR than in gap task in both species (**Fig. 3A**) (Brown et al. 2004). Right after the brief appearance of the target, a distinct peak of saccades was observed (component 1) (**Supplementary Fig. 1**, after the cyan line) and the saccades were mainly directed towards the target location in marmosets (79%, **Fig. 2E, 6A**). Although we did not observe this result in our human subjects, this result is similar to other research that explored humans’ express saccade makers. They showed short latency (around 80 to 120 ms SRT) express saccades when performing gap task, but they were unable to suppress visually evoked saccades, or “brief glancing” as described in the previous report, while doing a visual memory task (Fischer and Ramsperger 1984; Biscaldi et al. 1996). This brief glancing is usually attributed to a weaker voluntary control over saccade generation caused by a weaker fixation system either in the cortical or the subcortical areas such as the dorsomedial frontal cortex or the SC respectively (Bon and Lucchetti 1992; Hafed et al. 2009; Munoz and Wurtz 1993).

The second component (component 2) appears in delay periods of more than 200 ms in ODR task and seems to be more time locked with fixation offset in both marmosets and humans (**Supplementary Fig. 1**, before the magenta line), indicating that the prediction of the fixation offset contributes to the higher predictability, causing short SRT (Heeman et al. 2019). This is referred to as the urgency to respond with a premature saccade before the go cue (Brown et al. 2004). Consistent with previous descriptions, these saccades were partially directed toward the target (marmosets: 50.9%, humans: 66.7%, **Supplementary Fig. 3**). For the ones that were not directed toward the target, we suspect that it is a counter-balance effect of programming an antisaccade to avoid making a saccade after the target onset (Tian et al. 2018). Combining the two components, we observed an increase in impulsive error frequency and a prolongation of impulsive SRT with a longer delay period in both species (**Fig. 4, Fig. 5**).

As for the impulsive error in gap task, we found that they were all the so-called “anticipatory saccades” as previously reported (Chen et al. 2021; Edelman and Keller 1996; Mayfrank et al. 1986). In both species, the anticipatory saccades frequency increased and the SRT prolonged with a longer gap period (**Fig. 4, Fig. 5**), and showed no correlation with target eccentricity (**Supplementary Fig. 3, Supplementary Fig. 5**). These results are consistent with the previous interpretation that anticipatory saccades are the pre-target activities somehow crossing a theoretical saccade triggering threshold in the FEF or the SC (Dias and Bruce 1994; Dorris and Munoz 1998; Munoz et al. 2000). Our results further propose that these activities are very sensitive to the temporal (**Fig. 4, Fig. 5**) but less sensitive to the spatial predictability (**Fig. 2C, Supplementary Fig. 3, Supplementary Fig. 5**). Furthermore, because the visual target information was not yet provided, not all the saccades were directed toward the target (**Fig. 6A)**, similar to one previous report (Kalesnykas and Hallett 1987).

### Visuomotor (VM) Error

VM error saccades directed towards the target stimulus failed to reach the target because of either saccade spatial error (hypometric or hypermetric saccade), temporal error (failure to release fixation after the go cue or insufficient fixation (gaze left the eye window before the time allowed so). In gap task, saccade undershooting increased with target eccentricity in both species (**Fig. 6C**), which is consistent with previously reported results in humans (Frost and Pöppel 1976; Gillen et al. 2013), however, this correlation was more prominent in marmosets. This may be due to the more restricted eye mobility of the marmosets, as revealed by their smaller oculomotor range (marmosets, ∼10°; macaques, 40–50°; humans ∼55°) (Freedman 2008; Guitton and Volle 1987; Pandey et al. 2020; Mitchell et al. 2014), making the undershooting more apparent than in humans. On the other hand, an increase in saccade undershooting accompanied by a decrease in saccade overshooting in ODR task in marmosets **(Fig. 6D)** resembles what has been reported in humans (Dafoe et al. 2007; Evdokimidis et al. 2006; Gillen and Heath 2014). This seems to be reminiscent of the saccadic range effect previously reported in humans (Kapoula 1985), with the caveat that the target eccentricity varies within a single session, which was different from our task design. However, other reports failed to report similar range effect results despite the similar task design to Kapoula (Findlay 1982; Gillen et al. 2013; Nuthmann et al. 2016). Further experiments are necessary to determine whether what we observed here is related to the range. On the other hand, previous research showed that memory-guided saccades are usually more hypometric than visually guided saccades (Nuthmann et al. 2016), which agrees with our data from both marmosets and humans (**Fig. 6C, 6D**).

Another interesting result seen in marmosets was the substantial amount of VM error trials with saccades directed away from the target in both gap and ODR tasks (**Fig. 6B**), or what is referred to as targeting error (Parẻ and Munoz 2001). With further analysis, we found that the marmosets did not simply repeat their previous answer (saccade direction) every time, but only did so when consecutive VM error trials occurred (**Fig. 6E, 6F**). This perhaps mimics the so-called “recurring mistakes” effect (Kuvaldina et al. 2019) in which the same mistakes were repetitively made after the first one. This is mainly described in humans when performing simple, repetitive tasks. Many research studied this phenomenon and some explained that recurring errors may happen when we make a conscious effort or repeat the task several times (Dienes and Scott 2005), others mentioned that the choice to ignore some information might be implicitly remembered so that for the next event, the same information will be used/executed (Allakhverdov 1993, 2000). The neural pathway of recurring mistakes is still unclear, but one brain region candidate for mediating this phenomenon could be the parietal cortex (Purcell and Kiani 2016). Our experiment suggests that the marmosets might be useful to identify the neural bases of this psychological phenomenon.

### Successful Trials

In gap task, SRTs shortening with a longer gap period comes in agreement with the gap effect (Saslow, 1967; Reulen, 1984; Tam and Ono, 1994; Pratt et al., 1999; Schiller et al. 1987). However, SRT shortening with a longer delay period in ODR task was never reported before (**Fig. 5**, right column). Based on our data, we think this was due to the ultra-fast saccades right after fixation offset, observed in both species (**Supplementary Fig. 1**, the third component, **Supplementary Fig. 3**). The subjects were likely anticipating the fixation offset because we fixed the delay period for each session. Interestingly, human subjects showed a similar shortening in SRT and discrete components in the SRT distribution (after fixation offset), which might indicate a similar neural mechanism underlining the temporal execution of successful saccades. Anticipation is driven by previous knowledge, prediction, strategy, beliefs or expectations rather than visual inputs (Badler 2006; Kalesnykas and Hallett 1987; Smit and Van Gisbergen 1989), and the brief target presentation, as well as the blocked task design, might have facilitated anticipatory saccades to occur. Despite the tasks differences, one candidate area to consider regarding prediction is the FEF, as previous studies have shown that internal representations of invisible target motion in FEF might be linked to predictive eye movements (Barborica and Ferrera 2003; Xiao et al. 2006).

### Saccade Peak Velocity Changes When Error Happens

The main sequence relationship is believed to reflect the optimization of the trade-off between saccade duration and accuracy (Harris and Wolpert 2006). It is commonly known that there is a fixed relationship between the saccade amplitude to the peak velocity and duration (Fuchs 1967; Bahill et al. 1975). But when comparing visually guided and memory-guided saccades in macaques, it has been shown that saccades made in response to remembered locations had lower peak velocities and longer durations than saccades made in response to a visible stimulus (Gnadt and Andersen, 1988; Powers et al., 2013). In our results (**Fig. 7**), we found that the previous statement is true for both marmosets (higher slopes in gap task) and humans (similar slope but higher exponent in gap task). Yet, what was striking was that even in a simple task like gap or ODR task, we found that successful saccades showed the highest peak velocities compared to saccades from error trials in both marmosets and humans, which is usually more related to choice confidence in decision-making tasks (Seideman et al. 2018).

Moreover, it has been recently noticed that the intrinsic value of the visual input comes with a substantial impact on the saccade motor commands, and that saccade peak velocity is not just dependent on saccade amplitude, but also on motivation, arousal, and inter and intra-individual variations (Xu-Wilson et al. 2009; Muhammed et al. 2020). In addition, for advanced motor preparation, the expected value comes with great significance and, various studies have shown that saccade velocity can increase if a high reward is anticipated (Takikawa et al. 2002; Chen et al. 2013; Di Stasi et al. 2011). More studies are needed to determine the possible cause of this change in peak velocity.

## CONCLUDING REMARKS

The current study proposed a generalizable way of synchronising and analysing error trials with successful trials in different visual tasks. Marmosets are a promising experimental candidate because they capture humans’ success and error saccadic properties. Furthermore, analysis of success and error trials of behavioural data will help to understand the neural mechanisms behind mistakes and to better interpret the cognitive state of a subject performing a task. More importantly, it will help to set up the behavioural baseline measurements for future NHP models to study different brain disorders (Scott and Bourne 2022).

## DISCLOUSURE

The authors declare that the research was conducted in the absence of any commercial or financial relationships that could be construed as a potential conflict of interest.

## AUTHORS CONTRIBUTION

W.A. did the behavioural training, data collection, data analysis, and drafted the manuscript. C.C. guided the research and edited and revised the manuscript. H.O. performed the marmosets’ MRI imaging and surgeries. T.I. supervised the project and approved the final manuscript. All authors shaped the final version of the manuscript and approved the submitted version.

## ACKNOWLEDGMENT

We thank Professor Richard Veale for revising the English of this manuscript.

## FIGURE LEGENDS

**Supplementary Figure 1. Example saccade reaction times (SRTs) of impulsive error, visuomotor error and successful trials in ODR task, aligned on peripheral target onset, and the detection of SRT components**.

Four components were identified in the SRT distribution (illustration). The first component (1) showed up at around 120 ms after the target onset and appeared only in marmosets’ data, and represents the impulsive responses to the visual target. The second component (2) showed up in both species with delay periods of ≥ 250 ms. Because this component was time-locked to the fixation offset, we speculated that this component likely reflects the urgency to make a saccade. The third component (3) coincides with fixation offset with a delay period of ≥ 150 ms in marmosets and ≥ 200 ms in humans. This is likely reflecting the prediction of the end of the delay period, likely because of our blocked task design. The fourth component (4) represents the regular saccades responding to fixation offset. All the trials in ODR task belonging to marmoset S (A) and human H (B) are shown separated by the delay periods. For each sub-figure, the cyan line indicates target offset (67 ms) and the magenta line indicates fixation offset (end of the delay), the histogram (Y-axis on the left) indicates SRT frequency after target onset, and the orange curve (Y-axis on the right) indicates SRT cumulative probability after target onset, the orange curves indicate the smoothed SRT data and the blue stars represent the troughs that were detected by applying the reversed peak detection. Illustration figures were replotted from the delay period of 350 ms. As shown in the illustration, four SRT components, two before and two after fixation offset, were identified (1-4). Note that there is a break in the SRT histogram and a discontinuity of cumulative curve around the magenta line for all delay periods >100 ms in marmoset S and >150 ms in human H, meaning that saccades before and after the magenta line are likely responding to different monitor event (target flash and fixation off, respectively). Also, note that the ultrafast SRT immediately after the fixation offset for delay >100 ms in marmoset S and delay >150 ms in human H indicates that the subjects are probably counting (predicting) the time to make a response.

**Supplementary Figure 2. Marmosets’ saccade landing positions in ODR task**.

Heat maps show the normalized probability distribution of the saccade landing positions combining both the visuomotor error (VM) and the successful trials, separated by target eccentricities (top row: 4°, bottom row: 6°) for each marmoset. Saccades with the target on the left visual hemifield were mirrored so that the right solid circle represents the eye window around the correct choice whereas the left dashed circle represents the eye window around the incorrect choice. In each figure, the highest probability is normalized to 1. Landing positions inside the solid circle are all from successful trials. The dashed circle indicates the mirror-image side used for statistics. A significant proportion of saccades landed inside the target eye window (solid circle) compared to saccades from visuomotor error trials landed within the opposite eye window (dashed circle), for both targets at 4° (p= 1.1×10^−9^, p= 9.2×10^−4^, and p= 4.2×10^−9^, for marmoset S, N and M) and 6° (p=3.9×10^−50^, p= 8.3×10^−20^, and p= 2.1×10^−35^, for marmoset S, N and M). Wilcoxon rank-sum test was used for statistical analysis.

**Supplementary Figure 3. Changes in success and error rates with the target eccentricities in marmosets and humans in both tasks**.

A) The trial proportion of impulsive and visuomotor (VM) error and successful trials against target eccentricities were plotted for marmosets separating gap (top row) and ODR task (bottom row) to see the correlation. Results of the Spearman linear correlation test used for the gap task are provided in each subfigure and the ones with significant correlation are marked with *. Wilcoxon signed-rank test was used for the ODR task and the significant correlation is marked with #. B) The same analyses were also performed on human data. We didn’t perform statistical tests on human errors due to the scarcity of error trials. But please note that a similar trend of increasing VM error with farther eccentricity in gap task was observed in both marmosets and humans.

**Supplementary figure 4. Boxplot of SRT for success and errors in gap and ODR tasks**.

In this figure, we plotted one example boxplot from each species, marmoset S (A) and human H (B), showing the SRT of the impulsive and the visuomotor (VM) errors and the successful trials from both gap and ODR tasks. Although the SRTs are usually not normally distributed, most of the papers still use mean SRT for statistical tests. Because the mean SRT is the easiest to understand and to do statistical analysis. We also used mean SRT for analysis but here, we provide the boxplots of the SRTs to show the distribution from the two example subjects. Please note that all the trends and correlations we mentioned in Fig. 8 with the mean SRT were still preserved with the boxplots and the median of SRTs.

**Supplementary Figure 5. Changes in saccade reaction time of success and error subgroups with the target eccentricity in marmosets and humans in both tasks**.

A) The saccade reaction time (SRT) of impulsive and visuomotor (VM) error and successful trials against target eccentricities were plotted for marmosets separating gap (top row) and ODR task (bottom row) to see the correlation. Results of the Spearman linear correlation test used for gap task are provided in each subfigure and the ones with significant correlation are marked with *. Wilcoxon signed-rank test was used for ODR task and the significant correlation is marked with #. B) The same analyses were also performed on human data. We didn’t perform statistical tests on human errors due to the scarcity of error trials. No significant correlation can be observed besides decreasing SRT with farther eccentricity in successful trials in marmosets.

