## Supplementary Figures for "Dissecting errors made in response to externally- and internally-driven visual tasks in the common marmosets and humans"

**Supplementary Fig. 1**

**A marmoset S**

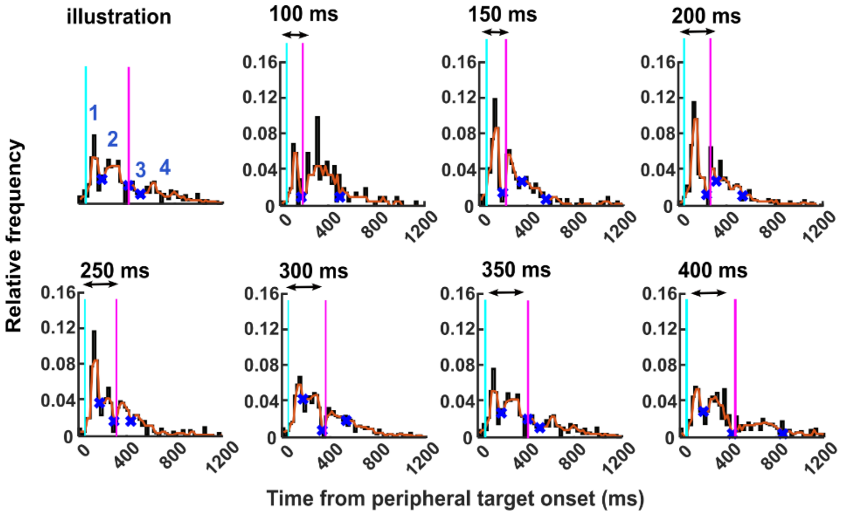


**B human H**

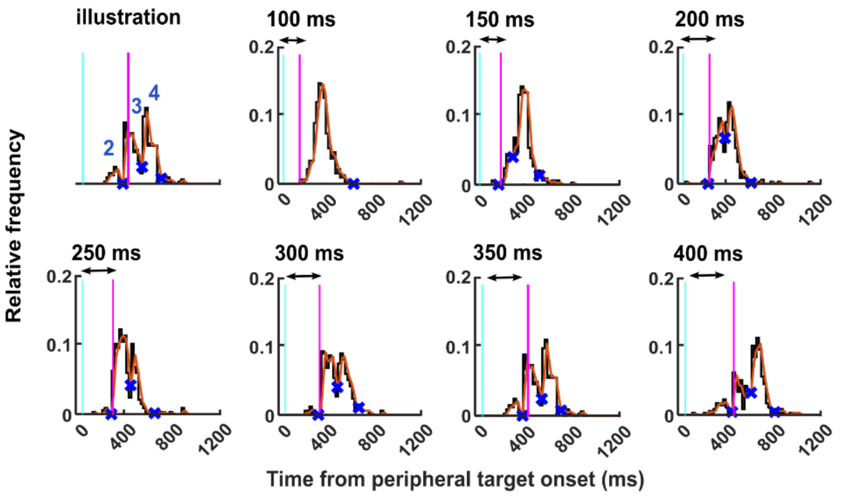

**Supplementary Fig. 2**

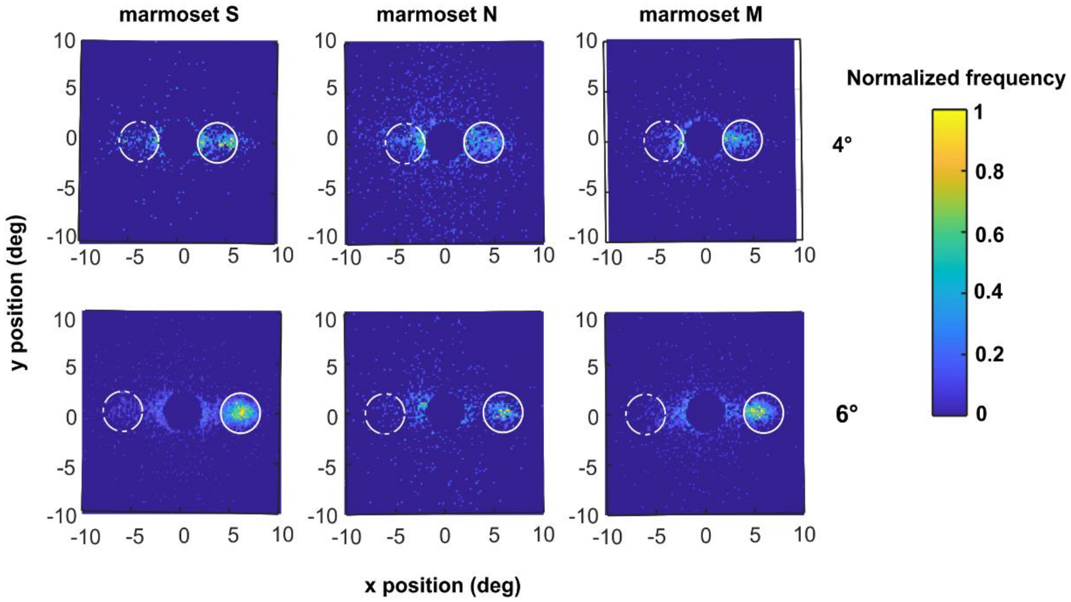

**Supplementary Fig. 3**

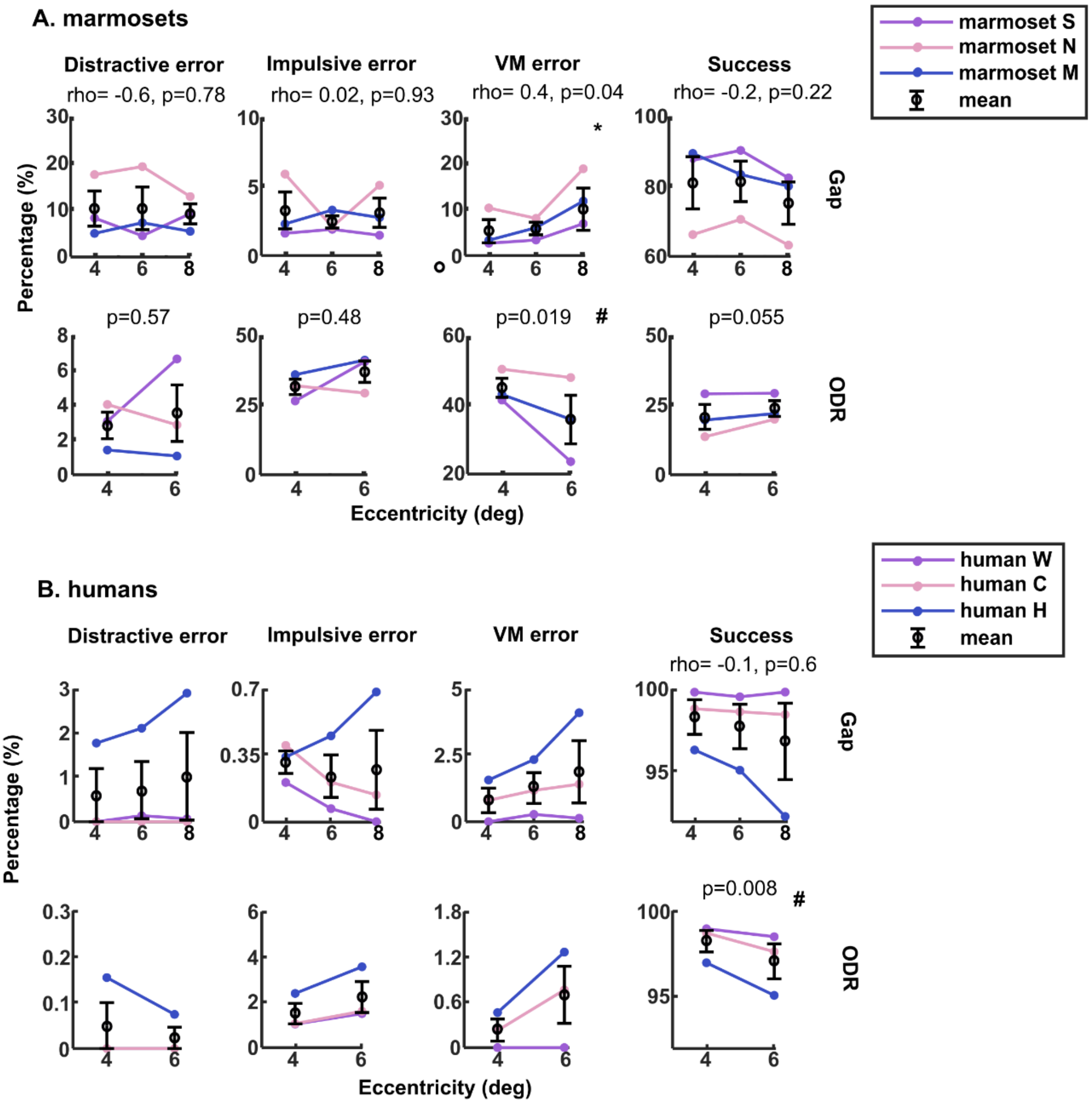

**Supplementary Fig. 4**

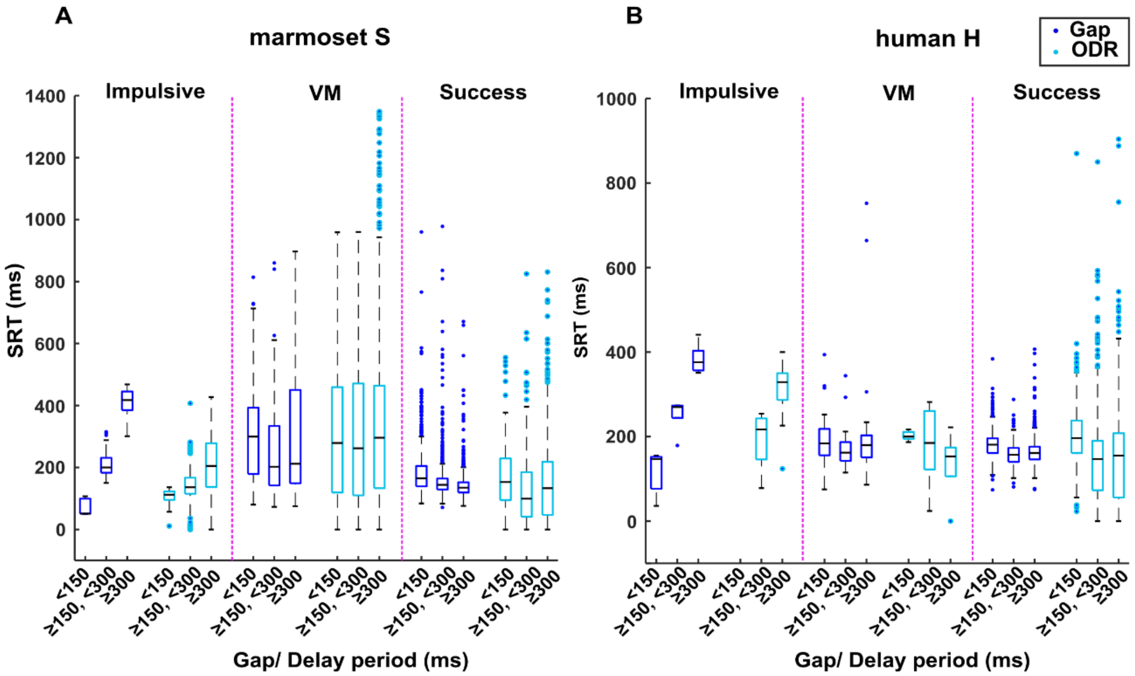

**Supplementary Fig. 5**

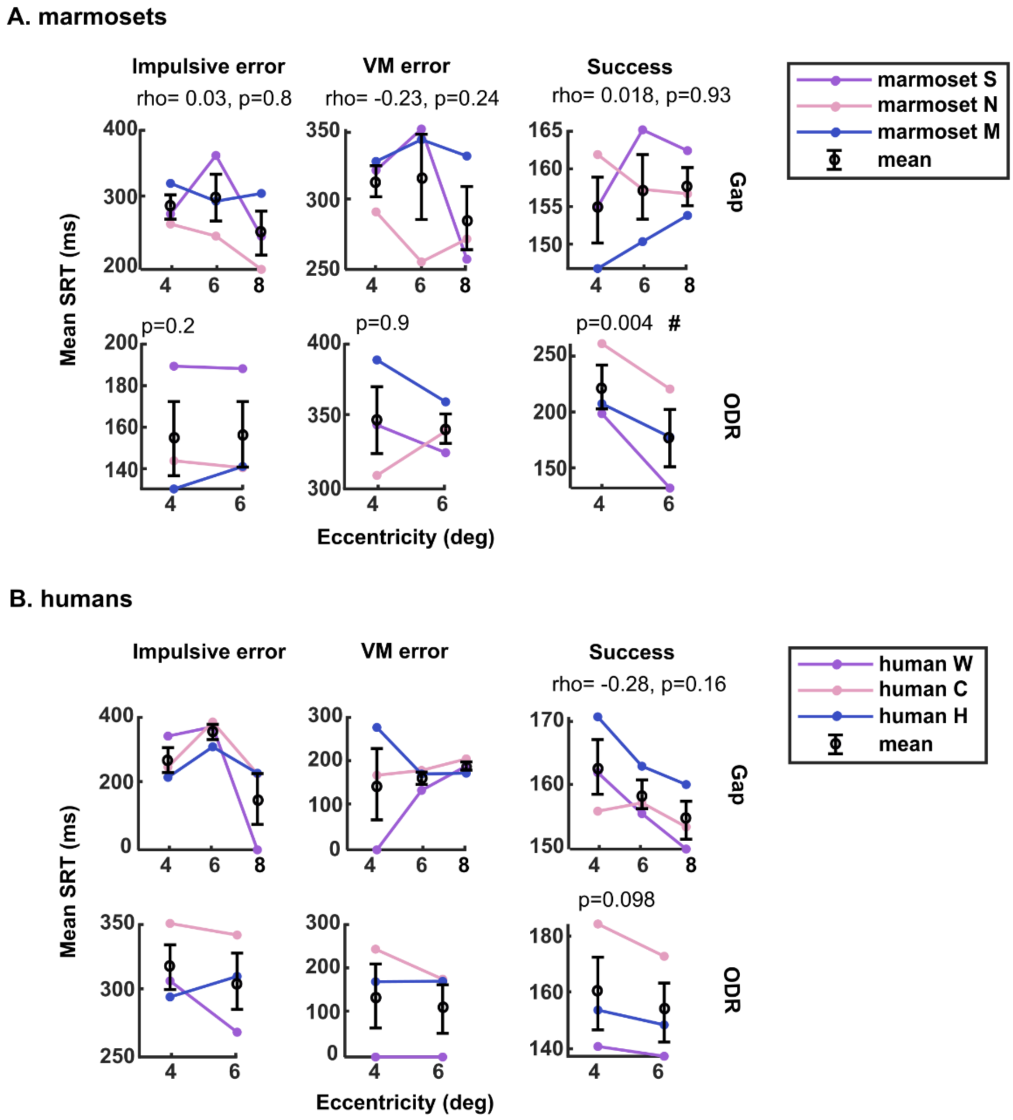
